# Prefrontal cortical control of a brainstem social behavior circuit

**DOI:** 10.1101/073734

**Authors:** Tamara B. Franklin, Bianca A. Silva, Zina Perova, Livia Marrone, Maria E. Masferrer, Yang Zhan, Angie Kaplan, Louise Greetham, Violaine Verrechia, Andreas Halman, Sara Pagella, Alexei L. Vyssotski, Anna Illarionova, Valery Grinevich, Tiago Branco, Cornelius T. Gross

**Author notes:** Current address: Sainsbury Wellcome Centre, University College London, UK.

## Abstract

The prefrontal cortex plays a critical role in adjusting an organism’s behavior to its environment. In particular, numerous studies have implicated the prefrontal cortex in the control of social behavior, but the neural circuits that mediate these effects remain unknown. Here we investigated behavioral adaptation to social defeat in mice and uncovered a critical contribution of neural projections from the medial prefrontal cortex to the dorsal periaqueductal grey, a brainstem area vital for defensive responses. Social defeat caused a weakening of functional connectivity between these two areas and selective inhibition of these projections mimicked the behavioral effects of social defeat. These findings define a specific neural projection by which the prefrontal cortex can control and adapt social behavior.

## Introduction

The medial prefrontal cortex (mPFC) plays an important role in generating appropriate social responses by supporting behavioral flexibility, response inhibition, attention and emotion. It has been proposed that the mPFC evaluates and interprets information within the context of past experiences, and is thus critical for selecting suitable behavioral responses within a social environment^1^. For example, lesions and pharmacological manipulations of the rodent mPFC modify inter-male aggression^2^, are required for sex differences in social anxiety^3^, modulate social position within a hierarchy^4^, and support the learned behavioral response to social defeat^5, 6^, highlighting the importance of this structure in interpreting and modifying social behaviors in the context of past social experiences.

The mPFC projects to several brain areas that are known to influence sociability, including amygdala, nucleus accumbens, hippocampus, and brainstem^7^. However, although several of these projections have been shown to be critical for mPFC control of non-social behaviors^8−10^, and mPFC projections to the raphe nucleus are able to interfere with the consolidation of adaptation to social defeat,^6^ until now the mPFC outputs that directly modulate social behavior have not been identified. Here we investigated whether projections from mPFC to the dorsal periaqueductal grey (PAG), a brainstem motor control area essential for defensive responses to social threats^11–13^, might play a role in the behavioral adaptation to social defeat. This adaptive response, occurring as a result of repeated exposure to threatening members of the same species, is characterized by a shift towards a more socially avoidant behavioral strategy^14^ that is presumably aimed at diminishing future harm and facilitating alternative routes to essential resources^15^. The adaptation to social defeat in animals may have clinical relevance because mood disorders, including major depression and social anxiety disorder, are thought to involve an extreme form of an adaptive coping strategy elicited by social adversity^16, 17^ ^18, 19^.

We found that repeated social defeat resulted in increased social avoidance and impaired working memory, both phenotypes that were ameliorated by the antidepressant ketamine. Selective pharmacogenetic inhibition of mPFC projections to PAG mimicked the effect of social defeat, increasing social avoidance and disinhibiting PAG. Social defeat caused a reduction in functional connectivity between mPFC and PAG, resembling observations made in imaging studies of patients with affective disorders^20^. Cell-type specific rabies virus tracing and *ex vivo* channelrhodopsin-assisted circuit mapping demonstrated that layer 5 mPFC projection neurons directly inhibit excitatory inputs to glutamatergic neurons in PAG and selective inhibition of these target neurons reduced social avoidance. These findings identify a specific projection by which the prefrontal cortex controls social behavior and demonstrates how these inputs can be modulated to adapt social behavior to the environment.

## Results

### Glutamatergic mPFC projections to dPAG

Anterograde and retrograde tracer studies have demonstrated prominent neural projections from the rat mPFC to the PAG^21, 22^. However, the precise location and cell identity of these projections have not been described. Moreover, although mPFC projection neurons are thought to be primarily glutamatergic, at least one study has demonstrated that GABAergic mPFC neurons project to the nucleus accumbens (NAc) and are capable of inducing avoidance behavior in a place-preference task^23^. To determine the identity of mPFC neurons that project to dorsal PAG (dPAG; we use this term to refer to the entire dorsal half of the PAG, including the dorsal-medial, dorsal-lateral and lateral columns), we simultaneously injected differentially fluorescent cholera toxin B retrograde tracers into NAc and dPAG (**Figure 1ab**) and visualized retrograde-labeled mPFC neurons. Labeled neurons projecting to NAc were located primarily in layer 2/3 with some labeled cells seen in layer 5 (**Figure 1c**). Labeled neurons projecting to dPAG, on the other hand, were exclusively located in layer 5 (**Figure 1d**), consistent with layer 5 harboring cortical projection neurons targeting brainstem motor areas^21, 22^. No overlap between NAc and dPAG projecting neurons was observed (0/791 and 0/594 neurons, respectively) arguing for a differential identity of these neurons in layer 5.

**Figure 1.**
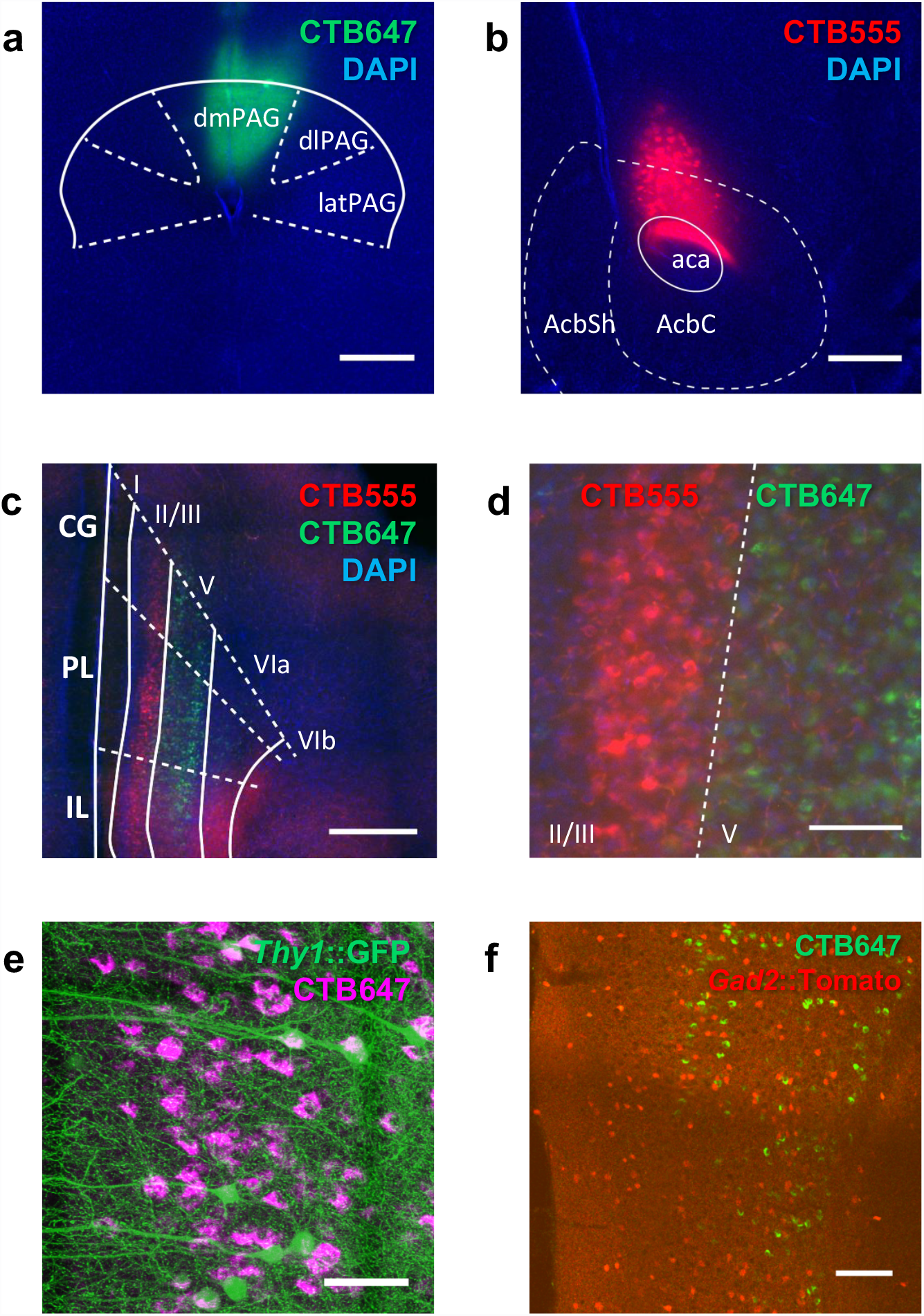
Layer 5 excitatory neurons in mPFC make direct projections to dPAG. (**a-d**) Mice were injected with retrograde tracers (CTB647, green) in dPAG (**a**) and (CTB 555, red) in NAc (**b**). Low (**c**) and high (**d**) magnification images of retrogradely labeled CTB647 (dPAG projecting) and CTB555 (NAc projecting) neurons in layer 5 and layer 2/3, respectively, of mPFC. (**e**) Representative image of retrogradely labeled CTB647 (dPAG projecting) neurons in mPFC of a *Thy1*::GFP mouse. (**f**) Representative image of retrogradely labeled CTB647 (dPAG projecting) cells demonstrating that these cells are not co-localized with GABAergic neurons in mPFC of *Gad2*::Cre;*RC*::LSL-Tomato mouse (scale bar = 500 µm in **a**-**c**, 100 µm in **d**, **f**; 50 µm in **e**). n=2.

To identify the specific cell-types involved, we first repeated the retrograde labeling experiment in *Thy1*::GFP-M transgenic mice^24^ in which sparse GFP labeling facilitates the morphological identification of neurons. Layer 5 mPFC neurons projecting to dPAG could be overwhelmingly identified as pyramidal in morphology, consistent with a glutamatergic identity (**Figure 1e**). Second, the retrograde labeling experiment was repeated in *Gad2*:: tomato transgenic mice in which GABAergic neurons are fluorescently labeled. No overlap between mPFC neurons projecting to dPAG and the GABAergic marker was detected (0/583 neurons; **Figure 1f**; **Supplementary Table 1**). These results suggest that, unlike the mPFC-NAc pathway, the mPFC-dPAG pathway consists exclusively of layer 5 glutamatergic projection neurons.

### Social defeat induces social avoidance

Chronic exposure of mice to an aggressor leads to social avoidance, but also causes more generalized changes in anxiety and depression-like behavior^25^ that might confound our search for plastic changes in the brain that drive behavioral adaptation to social threat. As a result, we sought to establish a sub-chronic social defeat paradigm associated with a selective adaptation of social behavior. Initially, we exposed male mice in their home cage once a day for five minutes to an aggressive conspecific confined behind a wire mesh barrier and then allowed them to freely interact for a further ten minutes, during which time the intruder repeatedly attacked the resident. Over seven days of social defeat, resident mice exhibited a gradual increase in upright submissive postures and freezing, and decrease in rearing during the direct encounter with the aggressor (**Figure 2a-c**). In addition, a gradual increase in social avoidance was observed during the anticipatory period in which the aggressor remained confined to the wire mesh barrier (**Supplementary Figure 1a**). Importantly, the number of attacks received by the resident did not differ across days (**Supplementary Figure 1b**) demonstrating that the changes in behavior elicited in the resident reflect a gradual adaptation to repeated social defeat. Because the behavioral adaptation of the resident tended to plateau after four days of social defeat we chose a three day defeat procedure for all further experiments to reduce potential generalization or habituation to the stress exposure.

**Figure 2.**
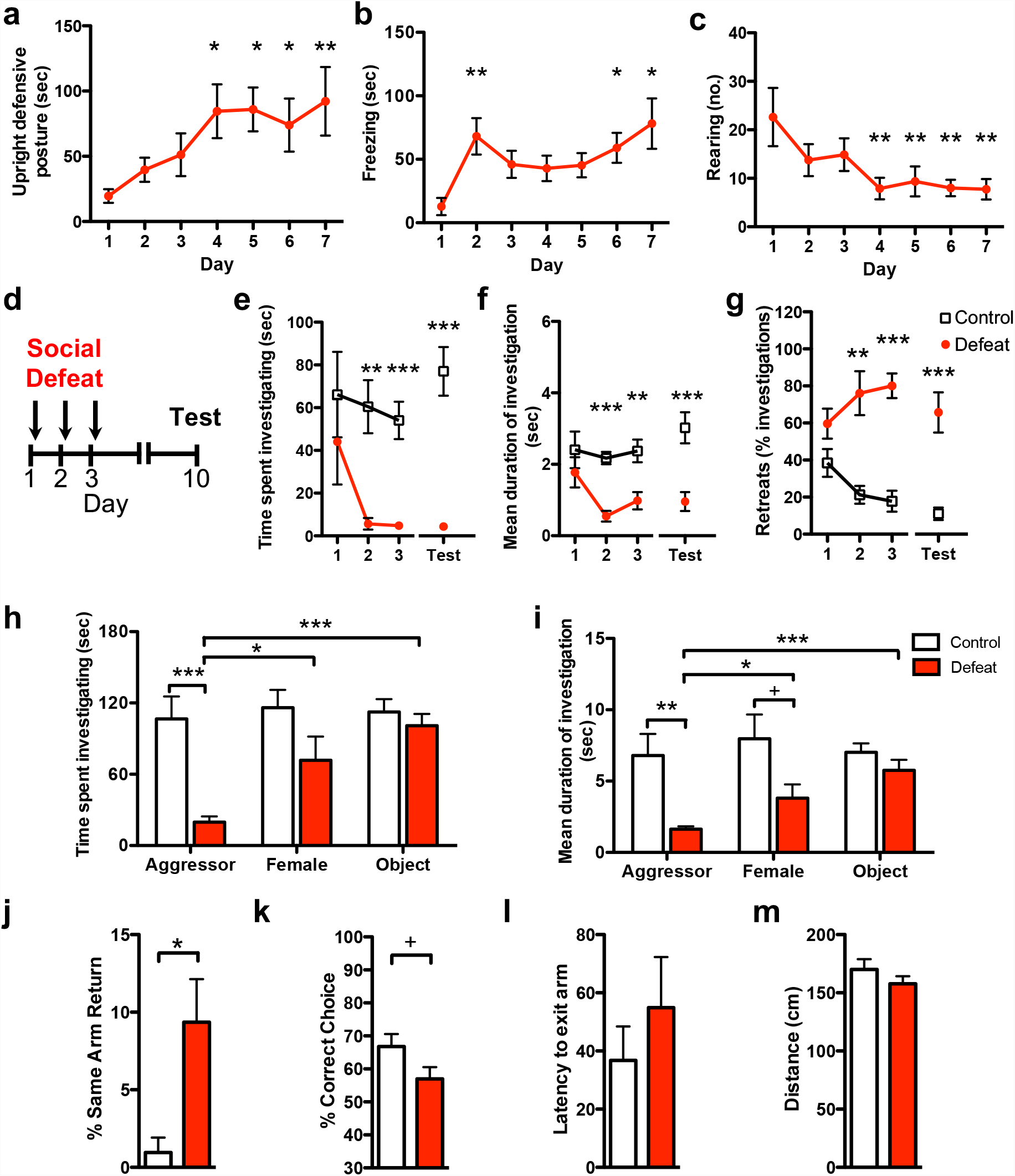
Induction of social avoidance by social defeat. Defensive responses elicited in the resident mouse by exposure to an aggressive intruder were increased across social defeat sessions as measured by significantly increased (**a**) upright-defensive postures (day: F_[6,7]_ = 3.8, P = 0.0042) and (**b**) freezing (day: F_[6,7]_ = 4.2, P = 0.0022), and decreased exploration as measured by (**c**) rearing (day: F_[6,7]_ = 3.2, P = 0.012). (**d**) Social approach behavior was measured each day for three days during an anticipatory period in which the intruder was restrained behind a wire mesh barrier immediately prior to social defeat or the control condition, as well as one week later (Test). Defeated mice (**e**) spent less time investigating a novel aggressor (defeat: F_[1,22]_=16.1, P = 0.006; day: F_[3,22]_ = 2.8, P = 0.047; defeat x day: F_[3,66]_ = 2.4, P = 0.079), (**f**) had shorter investigation bouts (defeat: F_[1,22]_=20.2, P=0.0002; day: F_[3,22]_ = 2.6, P=0.063, defeat x day: F_[3,66]_=2.1, P=0.11), and (**g**) retreated from social investigation periods more than control mice (defeat: F_[1,17]_ = 57.9, P < 0.0001; day: F_[3,22]_=1.9, P = 0.14; defeat x day: F_[3,51]_ = 8.7, P < 0.0001). All deficits persisted one week after the final defeat session. Defeated mice (**h**) spent less time (defeat: F_[1,12]_ = 7.6, P=0.018, stimulus: F_[2,12]_ = 12.4, P = 0.0002, defeat x stimulus: F_[2,24]_ = 8.9, P=0.0013) and (**i**) exhibited shorter investigation bouts (defeat: F_[1,12]_ = 7.5, P=0.018, stimulus: F_[2,12]_=5.0, P=0.016, defeat x stimulus: F_[2,24]_ = 3.9, P=0.033) toward both male and female intruders, but not a novel object when compared to control mice. In the Y-maze, defeated mice showed (**j**) increased same-arm returns (t_(14)_=2.9, P=0.013) and (**k**) a trend for decreased spontaneous alternation (t_(14)_=1.9, P=0.081), but (**l**) no change in latency to exit the start arm or (**m**) overall distance travelled. ^+^P<0.1; *P<0.05; *P<0.01; ***P<0.001). n=7-12.

To determine whether the sub-chronic social defeat procedure induced a persistent change in social coping strategy we monitored the behavior of the resident mouse during the anticipatory period immediately prior to each defeat session (Day 1-3), as well as during a test session (Test) in which an aggressor was placed into the resident’s cage within a wire mesh barrier one week later (**Figure 2d**). Resident mice spent progressively less time investigating the intruder both during the social defeat procedure and one week later (**Figure 2e**). Social defeat was accompanied by a progressive and persistent decrease in investigation bout duration (**Figure 2f**) as well as increase in the fraction of investigation bouts that were terminated by a rapid withdrawal movement, which we called “retreat,” **(Figure 2g**). Social defeat also elicited avoidance behavior when a female mouse, but not a novel object was placed into the wire mesh barrier on the test day, suggesting a selective adaptation of social behavior (**Figure 2hi**; **Supplementary Figure 1c**). In the Y-maze test, a short-term memory task known to depend on mPFC function^26^, defeated mice showed a significant increase in same arm returns, reflecting impaired working memory, but had normal latency to exit the arms and distance travelled, confirming unaltered exploratory behavior (**Figure 2j-m**). No significant changes in anxiety or stress-related behavior was seen in the elevated plus maze (**Supplementary Figure 1d-f**) or tail suspension test (**Supplementary Figure 1g**) confirming a selective impact of our defeat procedure on social behavior.

### Reversal of social avoidance by antidepressant treatment

Major depression is associated with increased social withdrawal and deficits in working memory that can be reversed by antidepressant treatment^27, 28^. To test whether the behavioral effects of social defeat demonstrated here might share pharmacological substrates with clinical depression we tested the effect of the rapidly acting antidepressant ketamine, an NMDA receptor antagonist, in our social defeat paradigm. On the day following social defeat animals received a single systemic injection of either ketamine (2.5 or 5 mg/kg) or vehicle and social interaction with an aggressive intruder was investigated one week later (**Supplementary Figure 2a**). Ketamine treatment was associated with a dose-dependent increase of time spent investigating the intruder (**Supplementary Figure 2b**). Ketamine did not significantly increase the duration of investigation bouts (**Supplementary Figure 2c**), but was associated with a dose-dependent reversal of the increased retreats induced by social defeat (**Supplementary Figure 2d**). No difference in locomotor activity was detected between control and ketamine-treated mice (**Supplementary Figure 2e**) suggesting a selective effect of the drug on social behavior. Ketamine treatment also ameliorated defeat-induced deficits in working memory, but had no significant effect on latency to exit the arms or distance traveled (**Supplementary Figure 2f-i**). These findings demonstrate that the persistent changes in social and cognitive behavior induced by sub-chronic social defeat depend on neural substrates shared with antidepressant treatment.

### Inhibition of mPFC-PAG projections mimics social defeat

To test whether mPFC-PAG projections might contribute to the behavioral effects of social defeat, we used a pharmacogenetic inhibition method to selectively suppress neurotransmission in mPFC-PAG projections. Mice were infected bilaterally in mPFC with adeno-associated virus (AAV) expressing the Venus fluorescent protein and HA-tagged hM4D (AAV-*Syn*::Venus-2A-HAhM4D)^13^, a designer Gα_i_-coupled receptor activated exclusively by the otherwise inert agonist clozapine-N-oxide (CNO)^29^, implanted with a guide cannula above the dPAG, and subsequently subjected to social defeat or control conditions (**Figure 3a, b**). Several weeks after infection HA-immunopositive afferents could be observed in PAG (**Figure 3c**) confirming the presence of hM4D on direct mPFC projections to this structure. CNO or vehicle was administered locally to the dPAG five minutes prior to behavioral testing one week after social defeat (**Figure 3d**). CNO-treated control mice spent less time investigating the aggressor, displayed shorter investigation bouts, and retreated more when compared to vehicle-treated control animals (**Figure 3e-g**, left). CNO-treated control mice were indistinguishable from vehicle-treated and CNO-treated defeated mice in time spent investigating the aggressor, duration of investigation bouts, and increase in retreats (**Figure 3e-g**), suggesting that mPFC promotes social interaction via direct projections to PAG. Additionally, social defeat may involve a weakening of mPFC-PAG projections, an interpretation that is consistent with the observation that CNO-treated defeated mice behaved similar to defeated mice administered vehicle (**Figure 3e-g**, right). CNO treatment did not affect overall locomotor activity arguing against a general role for these projections in exploratory behavior (**Supplementary Figure 3c**). Lastly, we performed a mPFC projection inhibition experiment where CNO was delivered to the overlying superior colliculus (SC), rather than the dPAG. In this experiment, no change in social interaction behavior was detected (**Supplementary Figure 3e-h**) suggesting that CNO delivery in the brain is local and affects a relatively restricted area.

**Figure 3.**
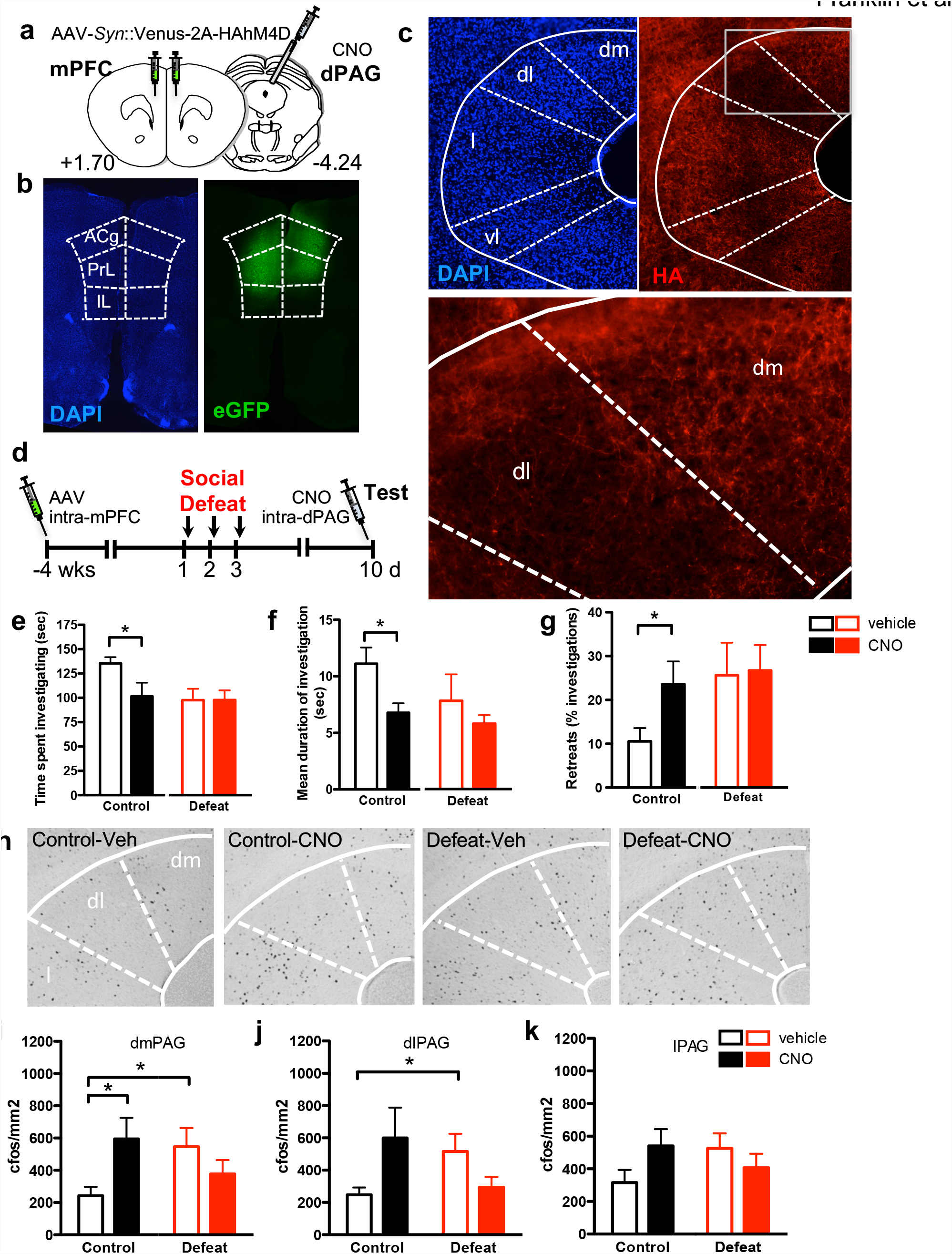
Inhibition of mPFC-dPAG projections mimics social defeat. (**a**) Mice were infected bilaterally in mPFC with AAV expressing Venus fluorescent protein and HA-tagged hM4D (AAV-*Syn*::Venus-2A-HA-hM4D), implanted with a guide cannula over dPAG, subjected to social defeat or control conditions, and infused locally in dPAG with CNO or vehicle before testing for social interaction. (**b**) Representative image of Venus labeled infected cells in the mPFC. (**c**) HA immunostaining revealed expression of hM4D in mPFC projections in the PAG. (**d**) AAV-*Syn*::Venus-2A-HAhM4D-WPRE was infused into the mPFC four weeks prior to social defeats. Social approach behavior was measured one week later (Test), immediately after intra-dPAG administration of CNO or vehicle. Control mice administered CNO prior to testing (**e**) spent less time investigating the aggressor (defeat: F_[1,1]_=3.54, P=0.067; CNO: F_[1,1]_=2.42, P=0.13; defeat x CNO: F_[1,39]_=2.32, P=0.14; t(19)=2.1, P=0.047) (**f**) exhibited shorter investigation bouts (defeat: F_[1, 1]_=2.23, P=0.14; CNO: F_[1,1]_=5.1, P=0.03; defeat x CNO: F_[1, 38]_=1.47, P=0.23; t(19)=2.9, p=0.0088) and (**g**) made more retreats (defeat: F_[1, 1]_=2.78, P=0.1; CNO: F_[1,1]_=0.54, P=0.47; defeat x CNO: F_[1, 38]_=2.5, P=0.12; t(19)=2.2, p=0.042), than vehicle treated control animals. Behavior of CNO-treated control animals was indistinguishable from vehicle-treated defeated mice and no effect of CNO treatment was detected in defeated animals. (**h**) Representative images and (**i-k**) quantification of cFos immunopositive cells in (**i**) dorsomedial (dm), (**j**) dorsolateral (dl), and (**k**) lateral (l) PAG of mice described above. Vehicle-treated defeated mice showed a significant increase in cFos immunopositive cells in dmPAG and dlPAG when compared to vehicle-treated control animals. CNO-treatment of control mice resulted in a significant increased in cFos immunopositive cells in dmPAG when compared to vehicle-treated control mice, matching levels seen in defeated mice (dmPAG, defeat x drug: F_[1,38]_=6.74, P=0.013, dlPAG, defeat x drug: F_[1,38]_=6.5, P=0.015). No significant effect of CNO treatment was observed in defeated mice. n=10-12. *P<0.05.

Following behavioral testing, animals were processed for cFos immunohistochemistry as an indirect measure of neural activity induced in dPAG by exposure to the aggressor (**Figure 3h-k**)^13, 30^. Vehicle-treated defeated mice showed significantly more cFos immunopositive neurons than similarly treated control mice in dPAG (dmPAG and dlPAG) suggesting that enhanced activation of dPAG is a neural correlate of social defeat and consistent with a role for this structure in defensive responses to a conspecific aggressor^11, 13^ (**Figure 3h-k** and **Supplementary Figure 3d**). CNO-treated control mice, on the other hand, showed a similar increase in cFos immunostaining across PAG subdivisions as socially defeated mice when compared to vehicle-treated controls (**Figure 3i-k**) demonstrating an inhibitory effect of mPFC inputs on PAG activity and corroborating a role for PAG in social avoidance. No further increase in the number of cFos immunopositive cells was seen in CNO-treated animals that had been exposed to social defeat when compared to similar vehicle-treated mice (**Figure 3i-k**) supporting the hypothesis that the effects of mPFC-PAG inhibition are occluded in defeated animals (**Figure 3e-g**).

### Social defeat weakens mPFC-dPAG functional connectivity

Deficient mPFC activity as well as reduced functional connectivity between mPFC and subcortical areas has been reported in persons experiencing major depression or social anxiety^31–35^ suggesting that mPFC-subcortical projections might be amenable to remodeling in response to social adversity. To determine whether social defeat might weaken mPFC-dPAG projections, we measured local field potential (LFP) coherence as a measure of functional connectivity between these structures in mice undergoing social defeat (**Figure 4a**). Social defeat was associated with a significant decrease in LFP coherence between mPFC and dPAG in both the theta and beta frequency bands in resident mice measured close to the intruder during the anticipatory period preceding social defeat when compared to control animals (**Figure 4b, c**). A similar trend was observed when the mice were far from the intruder (**Supplementary Figure 4b, c**). Moreover, Granger causality analysis of the LFP data revealed a shift in theta causality during defeat, with a significant increase in relative dPAG-mPFC causality found in defeated animals when compared to undefeated controls (**Figure 4d**, **Supplementary Figure 4d**). These results suggest a greater propensity for ascending information flow in this circuit following defeat. LFP spectral power in the theta band was decreased in defeated mice in both mPFC and dPAG relative to control animals (**Figure 4e-h**, **Supplementary Figure 4e-h**) suggesting that changes in oscillatory activity in one or both of these structures might underlie the altered functional connectivity in the theta frequency band. These findings are consistent with changes in LFP coherence in the theta frequency band reported between mPFC and both cortical and sub-cortical structures during cognitive and anxiety-related behaviors in mice that has been shown to reflect altered exchange or coordination of information between structures^36, 37^. Decreased coherence observed in defeated mice is not explained by any changes in oscillatory activity in either the mPFC or the dPAG (**Figure 4f, h**), suggesting a specific decrease in functional connectivity between these regions in this frequency band.

**Figure 4.**
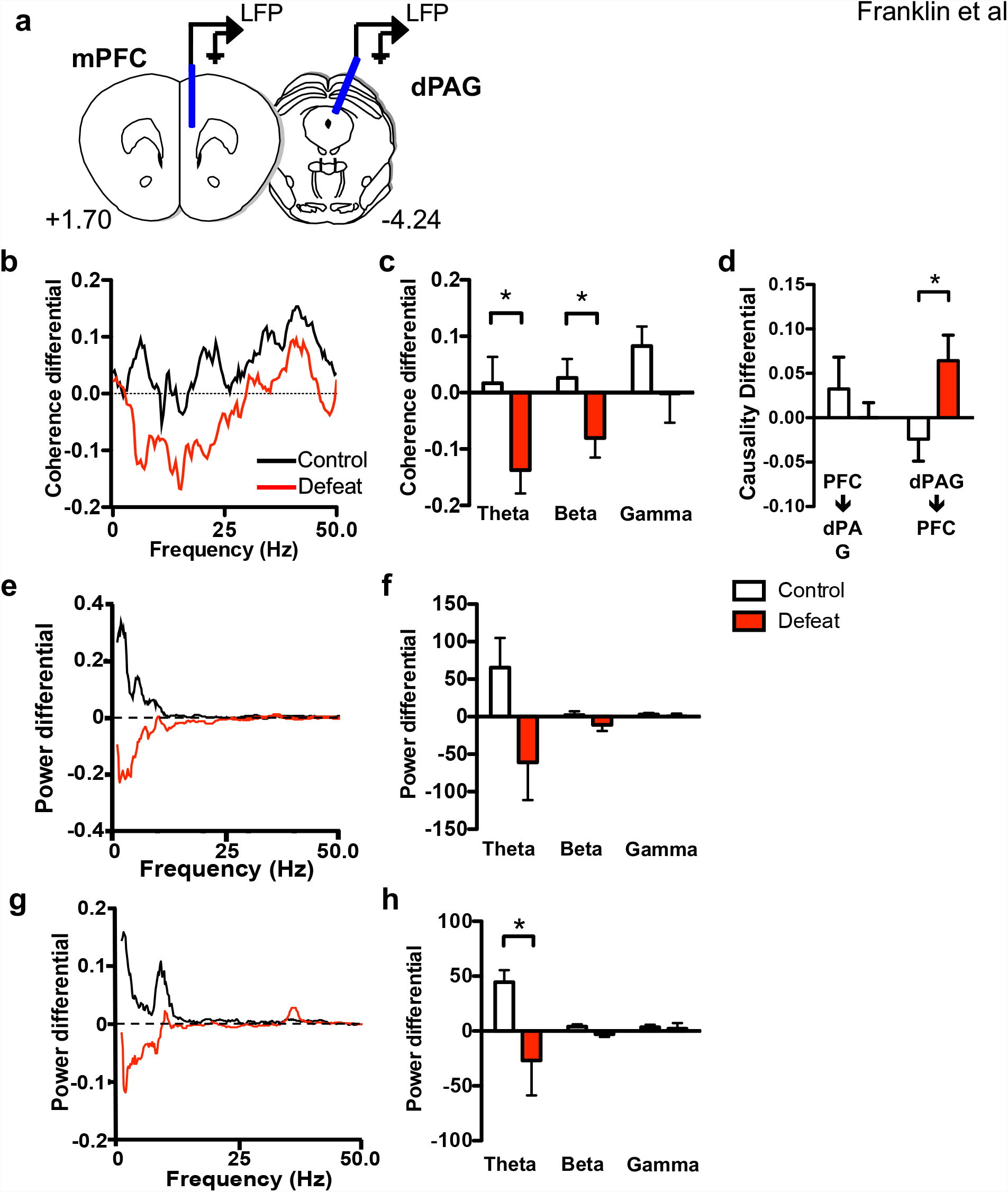
Social defeat weakens mPFC-dPAG functional connectivity. (**a**) Placement of electrodes used to measure local field potential (LFP) activity in mPFC and dPAG. Functional connectivity between mPFC and dPAG was estimated by measuring coherence between LFP signals at the two electrodes during the anticipatory period on day 3 compared to day 1 of social defeat. (**b, c**) Relative coherence (coherence differential) was significantly reduced in defeated mice compared to control animals (theta: U=9, p=0.048, beta: U=8, p=0.035). (**d**) Theta band causality between mPFC and dPAG was measured on day 3 compared to day 1 of social defeat. Relative causality (causality differential) was significantly higher in the PAG->mPFC direction in defeated mice compared to control animals (U=12, p=0.038). (**e-h**) Power spectra differential between day 1 and day 3 in (**e**, **f**) mPFC and (**g**, **h**) PAG when control and defeated mice were proximal to the aggressor. Defeated mice had lower power in the theta band in the PAG compared to control mice (U=6, P=0.018). Power spectra were averaged across mice. Power in each frequency band was calculated as the sum of the power values. n=7-8, *P<0.05.

Alterations in functional connectivity between brain structures as measured by LFP coherence can result from changes in synaptic connectivity between the structures, changes in the neural activity of one or the other structure, or changes in neural activity in a third structure mutually connected to the recorded structures. To test the first possibility we recorded evoked field potentials in dPAG in response to electrical stimulation of the mPFC in mice undergoing social defeat (**Figure 5a**). Periodic stimulation of mPFC during the habituation and barrier phases each testing day elicited short latency, multimodal population responses in dPAG (**Figure 5b** and **Supplementary Figure 5a**). No significant effect of social defeat could be detected across the experimental days on short latency response amplitudes (**Figure 5b**), despite significant avoidance developing in defeated animals (**Figure 5c**). However, changes in synaptic strength can be encoded either as changes in postsynaptic response amplitude or presynaptic release probability. To examine possible changes in presynaptic release probability in the mPFC-dPAG pathway during social defeat, we repeated the evoked LFP experiments using a double pulse protocol that allows for measurement of paired-pulse facilitation (PPF), a measure dependent on neurotransmitter release probability (**Figure 5d, e**). Initial experiments found maximal PPF in this pathway to occur at 50 ms pulse intervals (**Figure 5d**) and this interval was used for subsequent PPF monitoring. No significant differences in PPF were detected across testing days and groups (**Figure 5e**) suggesting an absence of synaptic plasticity in the direct mPFC-dPAG pathway during social defeat.

**Figure 5.**
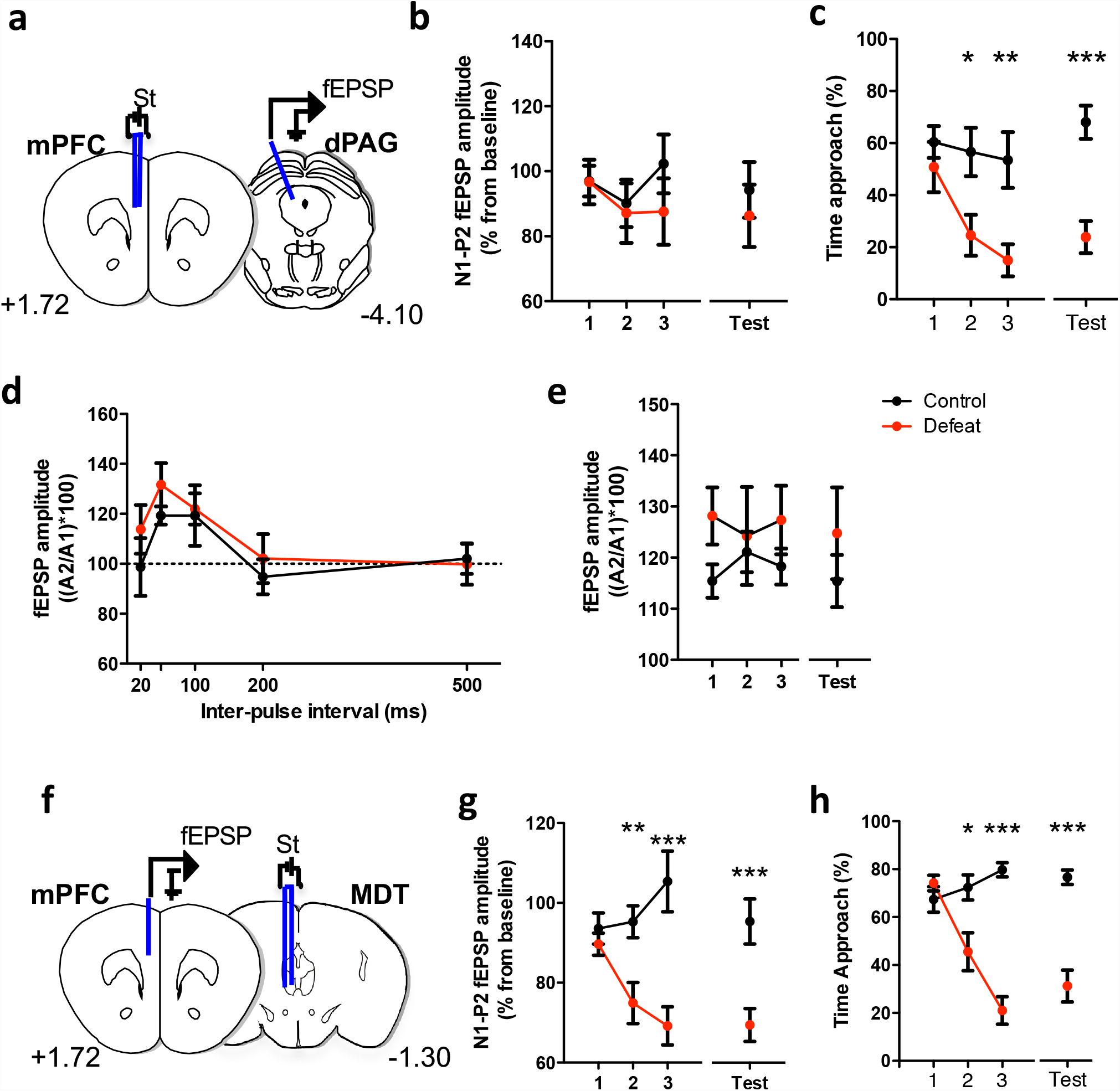
Evolution of synaptic field potentials in sensory and defeated mice across testing days. Location of recording and stimulating electrodes implanted chronically in the mPFC and dPAG (**a**)mPFC and MDT (**f**). (**b**) Similar mPFC-dPAG fEPSP amplitude in control and defeated mice but (**g**) significant difference in MDT-mPFC fEPSP amplitude in defeated mice compared to control (F_[3,51]_=5.58, p=0.0022). fEPSP amplitude is expressed as percent change in mean values (±SEM) during home cage exploration on the first day (baseline) for the N1-P2 interval during the social interaction (control: black circles; defeated group: red circles). Significant behavioral adaptation to social defeat in (**c**) mice with electrodes implanted in the mPFC and dPAG (F_[1,10]_=10.51, p=0.0088) (**h**) and mice with electrodes implanted in the MDT and dPAG (F_[3,57]_=13.93, p<0.0001). (**d**) Paired-pulse facilitation (PPF) of fEPSP recorded in the dPAG after stimulation of mPFC. Expressed as percent amplitude change (± SE) of the second fEPSP of the first for the five interpulse intervals. (**e**) Evolution of the paired-pulse facilitation of fEPSP along the sessions recorded in the dPAG after stimulation of mPFC. PPF, n=4; mPFC-dPAG, n=7; MDT-mPFC, n=10-12.

Next, we tested the possibility that reduced LFP connectivity between mPFC and dPAG could be driven by changes in afferent synaptic strength in mPFC. The mPFC receives prominent inputs from the medial dorsal nucleus of the thalamus (MDT) and reductions in this pathway have been hypothesized to occur in major depression^38^. To examine potential changes in this afferent pathway that could underlie weakened mPFC-dPAG functional connectivity we measured evoked field potentials in mPFC to stimulation of MDT during social defeat (**Figure 5f**). Periodic stimulation of MDT during the habituation and barrier phases each testing day elicited short latency, multimodal population responses in mPFC (**Figure 5g** and **Supplementary Figure 5b**). A significant reduction of short latency response amplitudes was detected across testing days in socially defeated mice when compared to control animals (**Figure 5g**), which paralleled the development of avoidance (**Figure 5h**).

These findings demonstrate that weakening of mPFC afferent synaptic strength occurs during social defeat and suggests that changes in mPFC afferent input strength underlies the weakened functional connectivity observed between mPFC and dPAG (**Figure 4b, c**).

### mPFC projections target glutamatergic dPAG cells

Our anatomical tract tracing (**Figure 1**) and pharmacogenetic projection inhibition (**Figure 3**) data argue that glutamatergic projections from mPFC act to inhibit dPAG function. To identify the local dPAG cell-types that mediate mPFC afferent control we performed cell type-specific monosynaptic circuit tracing using Cre-dependent pseudo-typed rabies virus^39^. Cre-dependent AAV expressing either the pseudo-typed rabies EnvA receptor TVA (AAV-*Ef1a*::DIO-TVA-mCherry) or the rabies virus protein G (AAV-*CAG*::DIO-RabiesG) were simultaneously delivered to dPAG of mice carrying either the *Vglut2*::Cre or *Gad2*::Cre transgenes^40, 41^ followed by infection with a pseudo-typed G-deleted rabies virus (∆G-EnvA rabies-GFP; **Figure 6a**). Following rabies infection brains were processed to systematically identify and visualize retrograde infected neurons (GFP+, mCherry-cells) across the entire brain rostral to the infection site. A total of 3231 cells were identified following infection of *Vglut2*::Cre mice (**Figure 6b-e**; **Supplementary Table 2**). The number of input cells present in each mouse was weighted to the density of starter cells in the dPAG at the centre of the infection site, and then averaged (**Figure 6e**). From the weighted averages, we observed that 90% of input cells were found in hypothalamus and thalamus, consistent with the major inputs of PAG deriving from diencephalic structures^42^. Only 6% of retrograde infected neurons resided in cortex, of which 20/182 were found in mPFC. Overwhelmingly, labeled mPFC neurons had a pyramidal morphology (**Figure 6c**) consistent with a layer 5 projection neuron identity (**Figure 1**)^43^. Similarly, 85% of cells identified following infection of *Gad2*::Cre mice resided in hypothalamus or thalamus (**Supplementary Figure 6a**; **Supplementary Table 3**), but we were unable to identify any retrograde labeled cells in cortex of infected *Gad2*::Cre mice. The relatively low frequency of long distance retrograde labeling in this line (total = 14 cells) suggested that long-distance afferents onto this class of cells are rare. These findings demonstrate that glutamatergic *Vglut2*^+^ neurons in dPAG are the major target of mPFC afferents and suggest that this cell class mediates the inhibitory input of mPFC on dPAG-mediated defensive responses.

**Figure 6.**
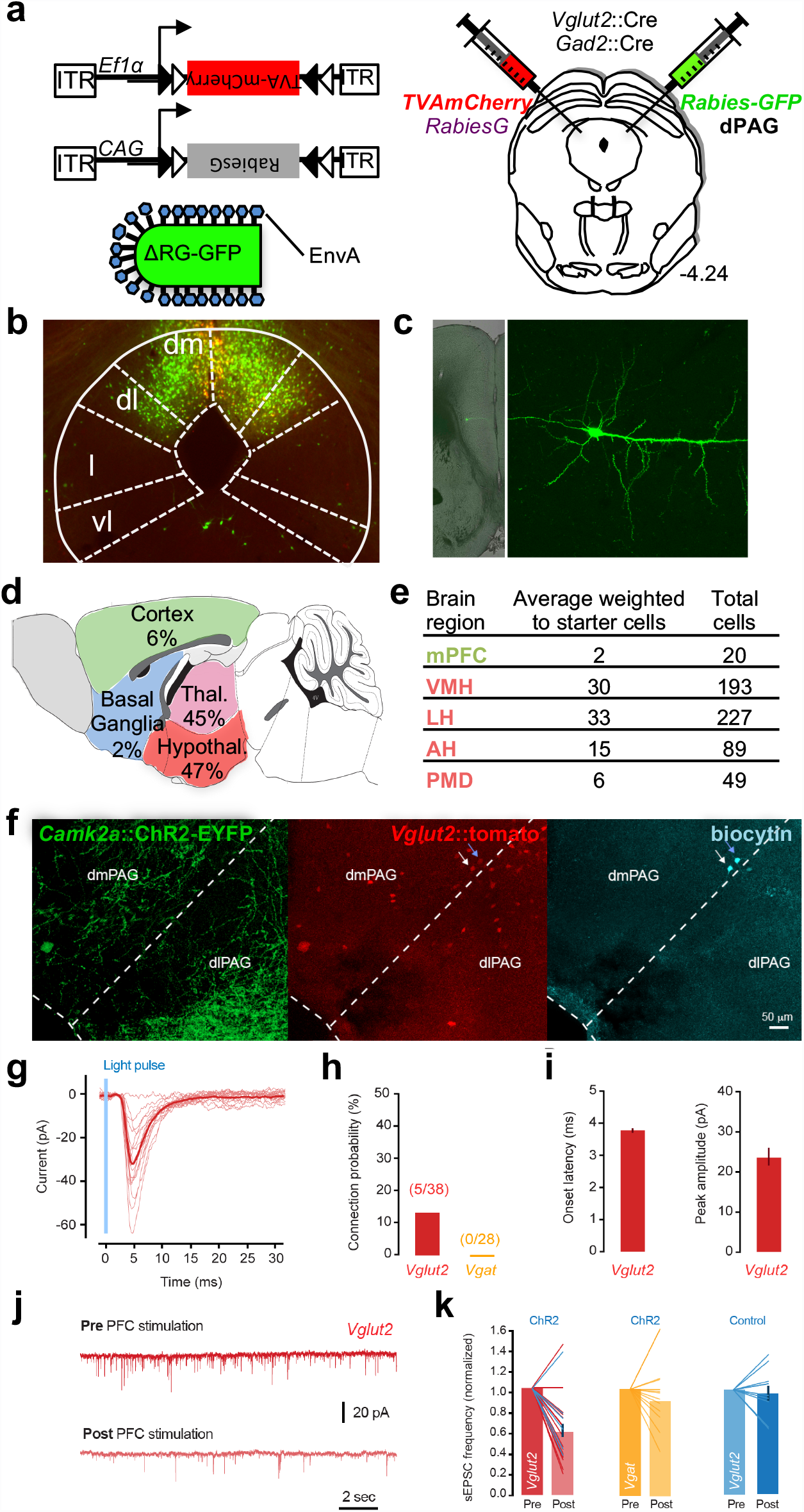
Cell-specific retrograde tracing identifies targets of PFC projections in PAG. **(a)** *Vglut2*::Cre and *Vgat*::Cre mice were infected in dPAG with Cre-dependent AAV expressing TVA-mCherry and rabies protein G and subsequently infected with EnvA pseudo-typed G-deleted rabies-GFP virus whose infection is limited to cells expressing TVA and that can form viable virions only in cells expressing protein G. In this manner infection by rabies-GFP is limited to cells expressing Cre and trans-synaptic infection occurs only monosynaptically. AAV and rabies were injected unilaterally into dPAG from opposing angles to avoid co-infection of the pipette tract. (**b**) Cre-dependent targeting of TVA-mCherry (red) and rabies-GFP (green) to Vglut2^+^ neurons in dPAG. (**c**) Low (left) and high (right) magnification images of a retrograde labeled rabies-GFP infected layer V pyramidal cell in mPFC. (**d**) Summary of rabies-infected neurons (GFP+, mCherry-) in the forebrain of *Vglut2*::Cre animals (percentage of the average number of retrograde neurons weighted to the number of starter cells present in each animal). (**e**) Number and weighted average of rabies-infected neurons in mPFC and hypothalamic nuclei (VMH, LH, AH and PMD) of *Vglut2*::Cre animals (n = 8). (**f)** Example images showing dense ChR2^+^ axonal projections (green) from the PFC in the PAG, cell bodies of Vglut2^+^ neurons (red) and two neurons filled with biocytin and processed after whole-cell recording (cyan). Blue arrow points to a neuron with monosynaptic input from the PFC and white arrow indicates a neuron without PFC input. (**g**) Light-evoked monosynaptic EPSCs in a Vglut2^*+*^ neuron. Light red traces are individual trials and dark red is average. (**h**) Mean probability of detecting PFC inputs in Vglut2 and Vgat neurons. (**i**) Average EPSC onset latency across all cells (left, 3.6±0.14 msec) and response peak amplitude (right, 23.1±4.5 pA). (**j**) Example traces of spontaneous EPSC recordings in a Vglut2^*+*^ neuron that did not receive direct PFC input, before and after 20 trials of ChR2^+^ stimulation (20 pulses at 10 Hz) of PFC terminals, showing a decrease in sEPSC frequency. (**k**) Mean change in sEPSC frequency with PFC ChR2 stimulation for all Vglut2^*+*^ neurons (left, 58.5±6% or control, P<0.0001, n=25) and Vgat^*+*^ neurons (middle, 91.4±8% of control, P=0.34, n=12). Right, light stimulation without ChR2 infection does not change sEPSC frequency. Lines show individual datapoints; in the left panel blue lines are cells with direct PFC input.

To test whether neural activity in *Vglut2*^+^ dPAG cells is selectively modulated by mPFC inputs as predicted by the rabies data, we performed *ex vivo* ChR2-mediated circuit mapping^44^. Following delivery of AAV-*CAG*:: ChR2-YFP to mPFC, acute slices were taken from dPAG and patch clamp recording was performed to examine light-evoked synaptic responses. Experiments were performed in either *Vglut2*::tomato or *Vgat*::tomato reporter mice to allow selective recording from identified glutamatergic and GABAergic neurons (**Figure 6f, Supplementary Figure 6b**)^45^. Short latency excitatory postsynaptic currents (**Figure 6g-i**) were identified in a small fraction (13%) of recorded *Vglut2*^+^ cells, but in none of the *Vgat*^+^ cells (**Figure 6h**). However, regardless of whether they received short latency inputs or not, the majority of *Vglut2*^+^ cells showed a significant reduction in the frequency of spontaneous excitatory inputs following ChR2 activation that was absent in control slices from non-infected animals (**Figure 6j-k**). *Vgat*^+^ cells, on the other hand, did not show a significant change in spontaneous excitatory inputs following ChR2 activation (**Figure 6k**) arguing for a selective inhibition of glutamatergic target cell afferents. Given the long latency of the inhibitory effect and the fact that the experiments were conducted under conditions in which light delivery failed to elicit action potentials, these findings demonstrate that glutamatergic mPFC projections directly suppress excitatory inputs onto *Vglut2*^+^ dPAG neurons via a presynaptic neuromodulatory mechanisms.

Finally, we examined the functional contribution of *Vglut2*^+^ and *Gad2*^+^ neurons in dPAG to social avoidance behavior during social defeat. Selective pharmacogenetic inhibition of neurons in dPAG was carried out by local infection of *Vglut2*::Cre or *Gad2*::Cre mice with AAV-*Syn*::DIO-hM4D-mCherry and subsequent systemic delivery of CNO 45 minutes prior to behavioral testing on day 10 (**Figure 7a**, **Supplementary Figure 7**). For *Vglut2*^+^ neurons a significant increase in time spent investigating the aggressor was seen in Cre+ mice when compared to Cre–littermates regardless of whether they experienced social defeat or not (**Figure 7b**). Inhibition of *Vglut2*^+^ PAG neurons had no significant effect on the duration of investigation bouts or the number of retreats (**Figure 7c, d**). No significant difference in avoidance behavior between Cre+ and Cre– mice was seen during the three days of social defeat prior to CNO administration (**Supplementary Figure 7a**) ruling out a confounding effect of genotype in these results. These data demonstrate that *Vglut2*^+^ neurons in dPAG are responsible for promoting avoidance during social interaction with an aggressor, a finding that is in line with the optogenetic activation of these cells producing defensive behavior and analgesia^46^ and the non-specific pharmacogenetic inhibition of this structure blocking defensive responses to social threat^13^. On the other hand, selective pharmacogenetic inhibition of *Gad2^+^* neurons elicited no significant change in time spent investigating the aggressor, nor duration of investigation bouts, although there was a decreased number of retreats in CNO treated mice when compared to vehicle treated littermates (**Supplementary Figure 7d-f**). These findings suggest that *Gad2^+^* neurons in dPAG do not make a significant contribution to social approach behaviour, at least under the conditions used in our experiments, but they may promote some aspects of defensive behavior.

**Figure 7.**
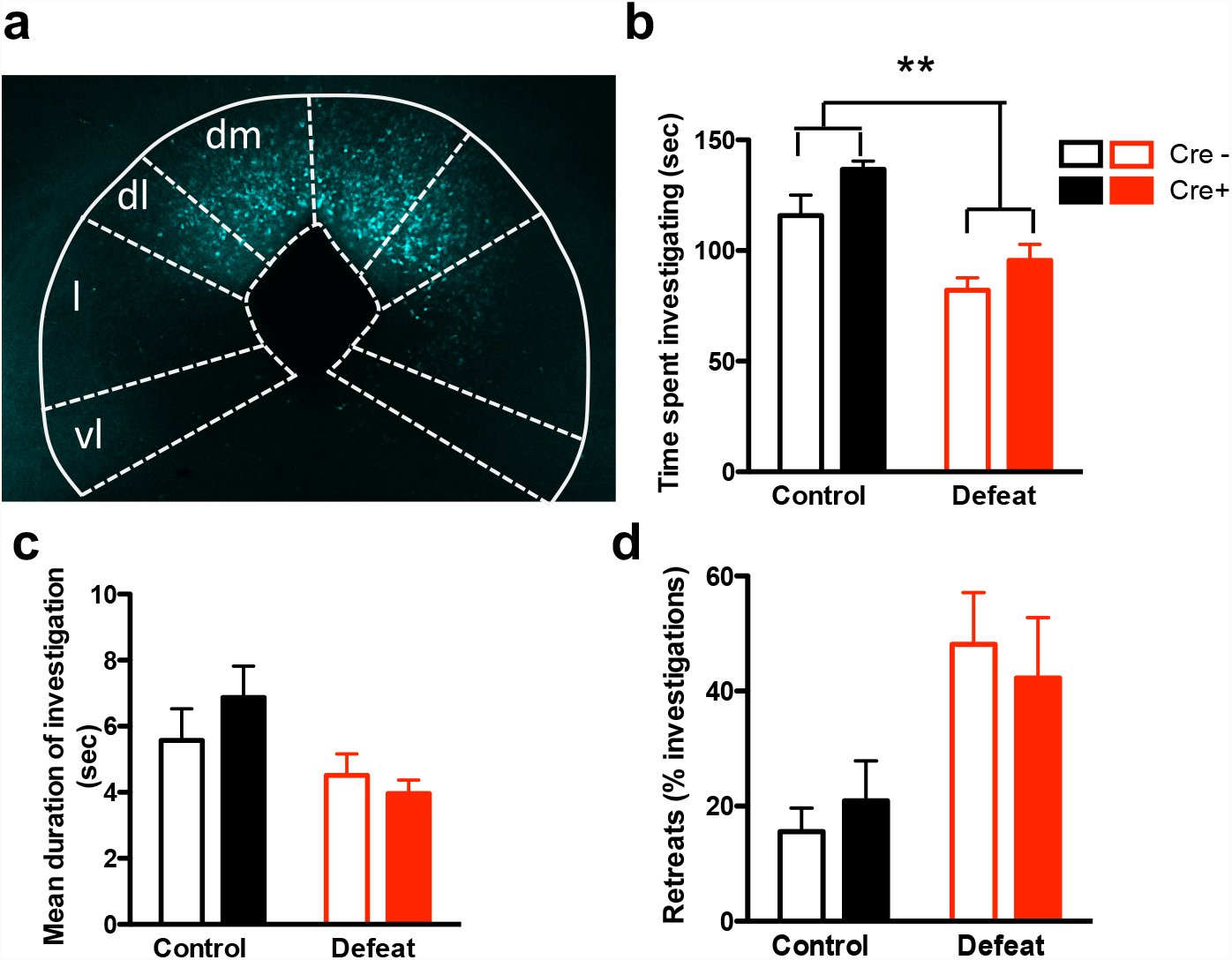
PAG inhibition increases social approach. (**a-d**) Selective hM4D-mediated inhibition of Vglut2+ neurons in dPAG. *Vglut2*::Cre mice were infected with AAV-*Syn*::DIO-hM4D-mCherry in the dPAG (**a**), subjected to control and social defeat, and treated with CNO before social interaction testing. Defeated mice (**b**) spent less time investigating the intruder (defeat: F_[1,11]_=26.77, P=0.0003), (**c**) had shorter investigation bouts (defeat: F_[1, 11]_ = 6.72, P = 0.025), and (**d**) made more retreats (defeat: F_[1,11]_ =22.28, P=0.0006) when compared to control animals. Systemic administration of CNO in Cre+ mice (**b**) increased time spent investigating the intruder (treatment: F_[1,11]_ = 13.12, P=0.004), but had no effect on (**c**) duration of investigation bouts or (**d**) retreats when compared to Cre-mice. n=6-7.

## Discussion

Considerable data has implicated neural activity in mPFC in the direct modulation of social behavior^4, 5^, but until now the projections mediating this effect were unknown. Our data demonstrate that the modulation of social approach/avoidance behavior by mPFC is mediated via direct projections to PAG, a structure required for the expression of innate motivated behaviors including defense, aggression, sex, maternal care, hunting, and foraging^47–51^. Moreover, the existence of major mPFC projections to both dorsal, defense-related, as well as lateral, approach-related, behavioral control columns in PAG (**Figure 3c**) suggests that these direct projections are likely to play important roles in the cortical modulation of behavioral adaptation under multiple environmental conditions, not just those described here. For example, firing of specific classes of neurons in mPFC has been shown to correlate with behavioral engagement and disengagement during foraging^52^ and mPFC is proposed to play a general role in decision-making in the face of environmental uncertainty.^53–55^

We used retrograde tracing, trans-synaptic rabies labeling, and *ex vivo* electrophysiology to show that layer 5 glutamatergic neurons in mPFC make monosynaptic excitatory connections onto glutamatergic neurons in dPAG and that, unlike mPFC projections to NAc, GABAergic neurons do not contribute to these afferents (**Figure 1** and **Figure 6**). Our discovery that these neurons are exclusively layer 5 excitatory pyramidal neurons is consistent with the known projections from this cortical layer to brainstem motor control areas involved in triggering and modulating behavior^21, 22^. Moreover, simultaneous retrograde labeling from dPAG and NAc showed that these mPFC projection neurons are non-overlapping (**Figure 1**). NAc afferents arise primarily from mPFC neurons residing in layer 2/3 and include long-range GABAergic neurons. This distinction suggests that different neuronal firing information is provided by mPFC to dPAG and NAc, a structure implicated in reward and behavioral selection. ChR2-assisted circuit mapping showed that only a small fraction (~10%) of *Vglut2*^+^ neurons in dPAG receive direct excitatory mPFC inputs, but that the vast majority receive strong indirect inhibitory mPFC inputs via a presynaptic neuromodulatory mechanisms (**Figure 6**). These findings suggest that glutamatergic mPFC projection neurons exert an inhibitory effect on dPAG by suppressing excitatory PAG afferents, possibly including those from medial hypothalamic regions that promote defensive behavior. Our findings raise the possibility that the small fraction of dPAG neurons receiving direct mPFC excitatory inputs may represent a specialized subclass of *Vglut2*^+^ neurons (**Figure 6**). Our behavioral findings showing that selective pharmacogenetic inhibition of *Vglut2*^+^, but not *Gad2*^+^ cells in dPAG increase social approach are consistent with a selective inhibitory presynaptic effect on *Vglut2*^+^ neurons in dPAG (**Figure 7 and Supplementary Figure 7**).

We established a sub-chronic social defeat procedure that induces a long-lasting increase in avoidance of social stimuli, including aggressive males as well as females, but not non-social stimuli, such as novel objects (**Figure 2**). Under these conditions, selective inhibition of mPFC-dPAG projections by pharmacogenetic hM4D-mediated projection inhibition caused a disinhibition of neural activity in dPAG and an increase in social avoidance (**Figure 3**). The observation that projection inhibition was not effective in socially defeated mice (**Figure 3**) suggested that the pathway was weakened by social defeat. This hypothesis was corroborated by LFP coherence data demonstrating a reduction of mPFC-dPAG functional connectivity in defeated mice and a switch in direction of causality with dPAG driving mPFC more strongly in defeated mice (**Figure 4**). Follow-up experiments using evoked field potential recording in behaving mice found that weakened functional connectivity between mPFC and dPAG was driven by a decrease in synaptic strength of afferent inputs to mPFC in the absence of any change in presynaptic or postsynaptic strength in the direct mPFC-dPAG pathway (**Figure 5**).

Our data have several implications. First, they support a critical role for dPAG in social behavior. Extensive lesion, pharmacological, and imaging data implicate dPAG in defensive responses to predators^56–59^. However, recent data show that dPAG is also required for flight, freezing, and avoidance behavior following exposure of rodents to aggressive conspecifics^13, 30^. Our findings extend this role to social avoidance in anticipation of threat (**Figure 2**). Such a role in modulating anticipatory avoidance is consistent with human imaging data demonstrating a rapid switch of BOLD signal activity from mPFC to dPAG in anticipation of predators^12^ or predator-like^59^ visual stimuli and suggests that dPAG may be involved in anxiety and as well as fear-related behaviors across species.

Second, our data demonstrate that functional connectivity between mPFC and dPAG can be moderated by social experience. Our *in vivo* evoked field potential experiments failed to find significant alterations in presynaptic or postsynaptic strength in the mPFC-dPAG pathway during defeat, but instead found a significant reduction in evoked responses in mPFC to thalamic stimulation (**Figure 5**). These data suggest that mPFC-dPAG functional connectivity is weakened by a reduction in upstream afferent drive during defeat. Numerous studies have found that dendrites of mPFC pyramidal neurons can atrophy in response to chronic stress^60, 61, 54, 55^ and reductions in the amplitude of excitatory inputs onto mPFC layer 5 pyramidal neurons were observed in subordinate mice and bidirectional manipulation of these receptors was sufficient to induce changes in stable hierarchies among cage mates^4^. Interestingly, one current theory of the physiological deficits underlying major depression proposes that reductions in thalamic inputs to mPFC are associated with a switch in mPFC processing from external to internal sensory information^38^.

While until now selective manipulation of mPFC outputs has not been shown to directly modulate social behavior,^6, 62^ Challis et al. (2014) has shown that mPFC-brainstem projections play a role in the induction of behavioral plasticity during social defeat. In this study, daily optogenetic activation or inhibition of mPFC terminals in the dorsal raphe nucleus immediately following social defeat blocked or precipitated social avoidance measured 24 hours after the last defeat experience. Because mPFC neurons provide excitatory input to local GABAergic neurons that tonically inhibit serotonin neuron firing in the raphe nucleus (and thus control serotonin release across the brain)^6^, mPFC projections may have a dual role in regulating global neuromodulatory tone (via dorsal raphe) and behavior (via dPAG) to achieve adaptation to social threats. It is, however, important to note that there are also key procedural differences between the current study and Challis et al. (2014). In our social defeat procedure, mice were tested for social avoidance in the same context as the aggression occurred, and thus our findings may be dependent to some degree on this aspect of classical contextual conditioning.

Both our cell-type specific retrograde rabies tracing and *ex vivo* electrophysiology experiments identified *Vglut2*^+^ neurons as the major target of mPFC projections in dPAG (**Figure 6**). Selective inhibition of *Vglut2*^+^ neurons in dPAG reduced social avoidance during presentation of the intruder (**Figure 7**) and recent studies have shown that optogenetic activation of this population of cells evokes defensive behaviors^46^. Our discovery that the vast majority of these cells receive presynaptic inhibitory inputs from mPFC provides a mechanism for the inhibitory effects of mPFC projections on cFos and social avoidance responses during exposure to an aggressor (**Figure 3**). The absence of either direct or presynaptic mPFC modulation of *Vgat*^+^ neurons (**Figure 6** and **Supplementary Figure 6**) and the absence of a behavioral effect of pharmacogenetic inhibition of this class of dPAG neurons was surprising, but suggests that cortical modulation of dPAG does not significantly depend on feedforward GABAergic inhibition.

Evidence from neuroimaging studies suggests that the mPFC-dPAG circuit we describe is likely relevant for understanding the prefrontal cortical control of human behavior. Direct projections between mPFC and dPAG have been described in primates^63^ and magnetic resonance imaging studies report a switch in brain activity from mPFC to dPAG during the pre-strike phase in a pseudo-predator video game situation^59^ suggesting that reciprocal activity in these structures may be involved in anticipatory fear in humans. While our study was limited to males due to its reliance on inter-male aggression, mPFC-dPAG projections are conserved across sexes and are likely to control instinctive behavioral outputs also in females. Electrical stimulation of human dPAG elicits the sensation of being chased, supporting its role in mediating avoidance responses to threat.^64^ Furthermore, our observation that the mPFC-dPAG-dependent social avoidance induced by social defeat can be reversed by treatment with a single dose of ketamine (**Supplementary Figure 2**), a potent antidepressant, suggests that this pathway may be a target of antidepressants that could serve as a neural substrate for the testing of antidepressant efficacy. Further work will be needed to identify the molecular mechanisms by which social experience remodels this pathway.

## Online Methods

### Animals

C57BL/6J and CD-1 mice were obtained from local EMBL or EMMA colonies, or Charles River Laboratories. CD-1 intruders were selected as aggressors if they attacked during the first 3 minutes after placement in the home cage of a novel C57BL/6J mouse across 3 consecutive days, as previously described.^65^ These mice typically represented the most aggressive 15% of CD-1 mice tested. *Vglut2*::Cre^40^ and *Gad2*::Cre (JAX stock 019022) mice were used in a heterozygous state. *Vglut2*::Cre;*RC*::LSL-tomato (called *Vglut2*::tomato), *Gad2*::Cre;*RC*::LSL-tomato (called *Gad2*::tomato), and *VGAT*::Cre; *RC*::LSL-tomato (called *VGat*-tomato) mice were obtained by crossing either the *Vglut2*::Cre, *Gad2*::Cre line, or *Vgat*::Cre line with *Rosa26*-CAG::loxP-STOP-loxP-tomato (JAX stock 007914). *Vglut2*::Cre;*RC*::LSL-EYFP (called *Vglut2*-EYFP) mice were obtained by crossing *VGlut2*::Cre (Jax stock 016963) with *Rosa26*-LSL-EYFP (Jax stock 006148). *Thy1*::GFP-M^24^ mice were used in a homozygous state. Mice were maintained in a temperature and humidity-controlled facility on a 12-hour light-dark cycle (lights on at 7:00) with food and water *ad libitum*. All behavioral testing occurred during the animals’ light cycle. All mice were handled according to protocols approved by the Italian Ministry of Health (#137/2011-B, #231/2011-B, #541/2015-PR) and commensurate with NIH guidelines for the ethical treatment of animals, except *in vitro* electrophysiology experiments which were conducted in the United Kingdom and were licensed under the United KingdomAnimals (Scientific Procedures) Act of 1986 following local ethical approval (Project Licence 70.7652).

### Social defeat

Singly-housed adult male mice (C57BL/6, 12-14 weeks old) were subjected to social defeat by placing an aggressive male CD-1 intruder mouse into the home cage of the experimental animal for 15 minutes each day. During the first 5 minutes the intruder was contained within a wire-mesh enclosure to prevent violent contact. Social approach and avoidance behavior, including number of investigations, investigation bout length, total time spent investigating, and number of retreats (sudden movement away from the intruder) was quantified during the first 3 minutes of this anticipatory period (Observer XT 11, Noldus) by an experimenter blind to the treatment group. For defeated mice, the wire-mesh enclosure was removed, after which the intruder invariably attacked the resident repeatedly. Submissive behaviors (freezing and upright defensive postures), and exploration (rearing) of the resident and aggressive attacks of the intruder were quantified during the ten-minute interaction period. Control animals were treated in the same manner, except that the wire mesh enclosure was not removed. This allowed control mice similar levels of visual, olfactory, and auditory contact with the aggressor as defeated mice.

### Social avoidance test

Five to seven days after the last social defeat session, animals were subjected to a social interaction test in which an aggressive CD-1 intruder (or a novel female or object, where specified) was constrained within a wire-mesh enclosure placed into the home cage of the experimental animal. The animals were allowed to interact through the wire-mesh barrier for 5 minutes, and approach and avoidance behaviors were scored in the same way as during the anticipatory period of social defeat. For mPFC-dPAG projection inhibition CNO was slowly infused via a single indwelling cannula (0.0015 mg, 0.15 μl, see below) immediately prior to testing. For *Vglut2*^+^ and *Gad2^+^* dPAG inhibition, all mice were first tested under control conditions, and then tested under defeat condition. Seven days after the last control session, and seven days after the last defeat session, CNO (3 mg/kg i.p.) or vehicle was systemically administered 45-60 min prior to testing. Testing consisted of a habituation session during which the experimental animal was allowed free exploration of their home cage for 5 minutes in the testing room, followed by the introduction of the intruder, behind a barrier, for a further 5 minutes.

### Y-maze

The Y-maze consisted of three grey, opaque plastic arms arranged at 120**°** angles around a center area. Animals were placed in a counterbalanced manner into one arm of the Y-maze and allowed to explore all arms of the maze for 8 minutes. Following a 2-minute habituation period, the percentage of correct choices and same arm returns were assessed for 6 minutes. A correct choice was quantified as each time the mouse entered all three arms without returning to an arm previously entered. Same arm returns (SARs) counted the number of times that a mouse entered fully into the center area and then returned to the arm they had just exited. Latency to exit the start arm and total distance travelled during the test were also quantified. Control and defeated mice were tested in the Y-maze one to two weeks after the last defeat session. Following the defeat treatment, mice either remained undisturbed, or were injected with vehicle, 2.5 mg/kg ketamine, or 5 mg/kg ketamine one day after the last defeat session. All injected mice tested in the Y-maze were also previously tested in the social avoidance test.

### Elevated Plus Maze

Mice were placed for 10 minutes on a four-arm plus maze made of two open and two closed arms (grey PVC, 30 cm x 6 cm) raised 50 cm above the ground. Manual scoring was done to quantify rearing and stretch attends in protected (body in closed arm) versus unprotected (body in open arm) areas as a measure of risk assessment. All elevated plus maze data was collected from surgeried mice previously tested in the social avoidance test.

### Tail Suspension Test

Mice were suspended by their tail from a hook (43 cm from floor) for 6 min. A plastic cylinder was placed around the tail to prevent tail climbing. All tail suspension data was collected from surgeried mice previously tested in the social avoidance test.

### Stereotactic surgery

Prior to surgery, mice were anesthetized with ketamine (100 mg/kg, i.p.) and xylazine (10 mg/kg, i.p.) and placed in a stereotactic frame (Kopf Instruments); isoflurane in oxygen was administered, as needed, to maintain anesthesia. For cholera toxin mediated retrograde tracing, the skull surface was exposed and mice were infused with 0.2 µl cholera toxin subunit B 0.5% (CTB647 and CTB555, Life Technologies) into dPAG (AP: −4.2; L: −1.18; DV: −2.36 from skull; angle: −26°) and into NAc (AP: +1.42 mm; L: −1.33 mm; DV: −3.5 mm from brain surface) using a glass capillary. In separate experiments *Thy1*::GFP (n = 8) or *Gad2*::Cre;*RC*::LSL-tomato (n = 1) mice were used. Serial coronal sections (250 *μ*m, except *Gad2*::Cre;*RC*::LSL-tomato, 50 μm,) were cut on a vibratome and visualized using confocal microscopy. For mPFC-dPAG or mPFC-SuColl projection inhibition, the skull surface was exposed and mice were infused bilaterally with 0.2 µl of an adeno-associated virus expressing Venus and hM4D (AAV-*Syn*::Venus-2A-HAhM4D-WPRE^13^) using a glass capillary filled with 1 µl of virus that was lowered unilaterally into the mPFC. After a 2-minute delay, the capillary was retracted, and the contralateral mPFC was similarly infused. For local CNO delivery a single 26-gauge stainless steel guide cannula (PlasticsOne) was implanted after viral infection into dPAG (AP: −4.16 mm; L: −1.0 mm; DV: −1.98 mm, angle: −26°; 1.25 mm projection from the pedestal), or into SuColl (AP: −4.1 mm; L: −0.75 mm; DV: −1.85 mm, angle: −30°, .20 mm projection from the pedestal) and secured to the skull using dental cement. For LFP recordings, the skull surface was exposed and two stainless steel watch screws were fixed permanently into the posterior and anterior portions of the skull, to serve as a ground and a reference, respectively. Teflon-coated tungsten wire electrodes were implanted unilaterally into PrL or Cg^66^ (AP: +1.65 mm; L: −0.50, DV: −1.50 mm from brain surface) and dPAG (AP: −4.16 mm; L: −1.32 mm, DV: −2.00 mm from brain surface, 26° lateral angle). Implanted electrodes were cemented directly to the skull with dental cement (DuraLay). For mPFC-dPAG and MDT-mPFC evoked potentials, animals were implanted unilaterally with bipolar stimulating electrodes into mPFC (AP: +1.72 mm, L: −0.40 mm, DV: −1.35 mm from brain surface) or MDT (AP: −1.2 mm, L: −0.40 mm, DV: −3.250 mm from brain surface) and a recording stereotrode into dPAG (AP: 4.1 mm, L: −1.3 mm, DV: −2.35 mm from skull surface, 26° lateral angle) or mPFC (AP: +1.72 mm, L: −0.40 mm, DV: −1.35 mm from brain surface) respectively. Electrodes were made of 50 μm, Teflon-coated tungsten wires (Advent Research Materials) and were used for stimulation or recording purposes as needed. A 0.1 mm bare silver wire was affixed to a stainless steel watch screw fixed permanently in the skull as a ground. The wires were connected to two three pins sockets (Archer connectors-M52). The connectors were fixed directly to the skull using acrylic resin (DuraLay) and connected to the Plexon system using a home-made adaptor. For rabies-mediated retrograde tracing, *Vglut2*::Cre and *Gad2*::Cre mice were infused into dPAG as described above with 0.1 µl AAV helper viruses that provided Cre-dependent expression of TVA and Rabies protein G (AAV-*EF1a*::DIO-TVA-mCherry-WPRE, AAV-*CAG*::DIO-RabiesG-WPRE; UNC Vector Core) followed 2-3 weeks later by infusion of an EnvA pseudo-typed rabies virus in which the protein G gene is replaced by GFP (1 µl; Salk Institute Vector Core^39^). AAV and rabies were both targeted towards the midline, but injected unilaterally on opposite sides to avoid co-infection of the pipette tract. For cell-specific inhibition in dPAG *Vglut2*::Cre or *Gad2*::Cre mice were infused 14 days prior to testing with 0.2 μl of AAV expressed hM4D in a Cre-dependent manner (AAV-*Syn*::DIO-hM4DmCherry-WPRE; UNC Vector Core). Serial coronal sections (70 μm) were cut on a vibratome and visualized under a microscope to verify placement of all electrodes, cannulas, and virus infections (**Supplementary Figure 3a, 4a**). Only mice with appropriate placements were included in the reported data. For *in vitro* electrophysiology, *Vglut2*::Cre;*RC*::LSL-tomato or *Vgat*::Cre;*RC*::LSL-tomato male mice were injected bilaterally into mPFC (AP: +1.7; ML: ±0.6; DV: −1.35) with 0.05 ul of AAV2-*CamKIIa*-hChR2(H134R)-EYFP virus (UNC Vector Core) delivered via manual hydraulic pump (Narishige). Following injection mice were allowed at least 2 weeks for viral expression.

### In vivo *electrophysiology*

All mice were allowed to recover for at least 7 days before testing and mice were habituated repeatedly for several days to the recording device by attaching a mock device of similar size and weight. LFP recordings were performed using a battery-powered custom wireless amplifier and recording device (23 x 15 x 13 mm, 3.7 g) located on the head of the animal^67, 68^. LFP signals from electrodes located in mPFC and dPAG were sampled at 1600 Hz (bandpass filter 1-700 Hz) and stored in the on-board 1 GB memory chip at 1600 Hz^69^. A built-in accelerometer registered the movements of the animal throughout the experiment and an infrared detector on the device was used to synchronize electrophysiological and video recordings. For evoked potential recordings, the neural signal was amplified (gain 1000x) and filtered (bandwidth of 0.1Hz-10 kHz) through a headstage and a differential pre-amplifier (Omniplex, Plexon). Signals were digitized at 40 kHz and continuous recordings were collected for offline analysis. Synaptic field potentials in dPAG were evoked using a pulse generator (CS-420, Cibertec) and electrical stimulator (ISU-200bip, Cibertec) during homecage exploration and while the intruder was present in the home cage behind a barrier using a single 100 μs, square, biphasic (negative-positive) pulse applied to mPFC at a rate of 0.1 Hz. For each animal, the stimulus intensity was 40-50% of the intensity necessary for evoking a maximum fEPSP. Evoked potentials were monitored using an oscilloscope (Tektronix). At completion of the experiment, mice were anesthetized using 2.5% Avertin (400 mg/kg, i.p.; Sigma-Aldrich) and perfused transcardially (4.0% wt/vol paraformaldehyde, 0.1M phosphate buffer, pH 7.4). For LFP recordings, a small electrolytic lesion was made around the tip of the electrode (0.4 mA, 3 s; Ugo Basile Lesion Making Device, Ugo Basile) before the animal was perfused. Serial coronal sections (40 or 70 μm) were cut on a vibratome and visualized under a microscope to verify all electrode placements (**Supplementary Figure 3**).

### In vitro *electrophysiology*

Acute coronal slices (200 μm, containing the PAG were prepared from 11-13 week old mice. Animals were killed by decapitation following isoflurane anaesthesia. Coronal slices were cut at 4°C using a 7000smz-2 vibrating microtome (Campden, UK). Brain slices were incubated at 37°C for one hour before being kept at room temperature prior to experiments in artificial cerebrospinal fluid (ACSF) containing: 125 mM NaCl_2_, 2.5 mM KCl, 26 mM NaHCO_3_, 1.25 mM NaH_2_PO_4_, 25 mM glucose, 2 mM CaCl_2_, 1 mM MgCl_2_, 0.2% biocytin (pH 7.3 when bubbled with 95%O_2_ and 5%CO_2_). Boroscillicate glass micropipettes with a 3-6MΩ resistance (Harvard Apparatus, UK) were filled with: 136 mM K-Gluconate, 4 mM KCl, 10 mM HEPES, 1 mM EGTA, 2 mM Na_2_ATP, 2 mM Mg_2_ATP, 0.5 mM Na_2_GTP, filtered (2 μm, prior to patching. Fluorescent cells were visualized on an upright Slicescope (Scientifica, UK) using a 60× objective and the relative coordinates of each neuron were recorded. Whole-cell patch clamp recordings were achieved at room temperature, using a HEKA 800 Amplifier (HEKA, Germany). Data was acquired at 25 kHz using custom software. Channelrhodopsin was activated with widefield 490 nm LED illumination (CoolLED; 1ms pulses). After electrophysiological recordings, slices were fixed in 4% paraformaldehyde for 15 minutes, incubated in blocking solution for 30 minutes containing 5% normal goat serum and 0.3 Triton X 100 in PBS, followed by primary antibody at 4°C overnight. The slices were then washed with 0.3% Triton X 100 in PBS (PBS-T) for 3x10 min, incubated with secondary antibody for 1h at room temperature, and after 2x10 min washes in PBS-T, they were incubated for 20 minutes in PBS-T with streptavidin to visualize biocytin-labelled neurons. After an additional 10 min wash in PBS, slices were mounted in Slow Fade mounting medium (Molecular Probes). All antibodies used were from Molecular Probes: chicken anti-GFP (1:1000), Alexa Fluor 488-conjugated goat anti-chicken IgG (1:1000), Alexa Fluor 635-conjugated streptavidin (1:500). Recorded slices with biocytin-filled neurons were imaged with 10x and 40x objectives on a Leica SP8 inverted confocal microscope (Leica). Deconvolution was performed using Huygens Software (Scientific Volume Imaging) and tiling of individual images was done in Fiji (Schindelin et. al., 2012).

### Immunohistochemistry and immunofluorescence

Immediately following social avoidance testing mice were returned to their housing room for 90 minutes, deeply anesthetized with Avertin (400 mg/kg, i.p.; Sigma-Aldrich), perfused transcardially (4.0% wt/vol paraformaldehyde, 0.1M phosphate buffer, pH 7.4) and the brain was removed and post-fixed overnight in 4.0% paraformaldehyde. The posterior half of the brain was cryoprotected (30% sucrose wt/vol, 0.1M PBS, pH 7.4) at 4°C overnight and flash frozen in isopentane. Coronal sections were taken with a sliding cryostat (40 μm; Leica Microsystems) and immunohistochemistry was performed. For cFos visualization floating sections were incubated with anti-cFos antibody (1:10,000, Ab-5; Calbiochem) for 72 hours at 4°C, after which the primary antiserum was localized using the avidin-biotin complex system (Vector Laboratories). Sections were incubated for 90 minutes at room temperature in a solution of biotinylated goat anti-rabbit (PK-6101, Vector Laboratories) and then incubated in an avidin-biotin horseradish peroxidase complex solution (ABC Elite Kit, Vector Laboratories) for 90 minutes at room temperature. The peroxidase complex was visualized by incubating slices for 5 minutes with a chromogenic solution consisting of 0.05% wt/vol 3,30-diaminobenzidine tetrahydrochloride (Sigma-Aldrich), 6 μg/ml glucose oxidase (Sigma-Aldrich), 0.4 mg/ml ammonium chloride in PBS, and then adding 2 mg/ml glucose to the solution. The reaction was stopped by extensive washing in PBS and sections were mounted, dehydrated and cover-slipped with quick mounting medium (Eukitt, Fluka Analytical). cFos immunopositive cells were counted using manual thresholding and automatic counting (ImageJ) in a section chosen randomly (Bregma −4.16) by an investigator blind to experimental treatment.

For visualization of HA-tagged hM4D, slices were mounted onto SuperPlus slides and allowed to dry. Slides were then boiled in citrate buffer (10 mM, pH 6.0) for 10 minutes and allowed to cool to room temperature, before being submerged in PBS containing 0.4% Triton-X (PBS-T) for 1 hour. They were then placed in blocking solution (1% BSA, 5% Normal Goat Serum in PBS-T) at room temperature for 1 hour, followed by incubation with a rabbit anti-HA mAb (C29F4, Catalog #3724, Cell Signaling) at 1:500 in blocking buffer. Slides were exposed to secondary antibody (Alexa Fluor 546-conjugated goat anti-rabbit IgG, Invitrogen) in blocking buffer at room temperature for 90 minutes. Slides were then exposed to 4’,6-diamidino-2-phylindole, dihydrochloride (DAPI, Molecular Probes) at 1:1,000 in PBS at room temperature for 20 minutes. Slices were washed extensively with PBS between incubations and following DAPI staining.

### Electrophysiology data analysis and code availability

LFP data were analyzed using Matlab (Mathworks) with the Chronux toolbox (coherencyc, http://chronux.org/^70^). To assess synchrony between LFP signals coherence was calculated with the multi-taper method, using a 200 ms window, time-bandwidth product (TW) of 5, and 9 tapers. The Granger causality used an order of 20 estimated by a bivariate autoregressive model. LFPs in the mPFC and dPAG were recorded on the 1st day and 3rd day of social defeat during the anticipatory period. fEPSP slopes were analyzed off-line using commercial computer programs (Spike2 and SIGAVG, Cambridge Electronic Design) using the same rate period.

### Statistical analysis

Data analysis was performed using Statview (SAS) or Sigmaplot, except *in vitro* electrophysiology data, which was analyzed in Python 2.7 using custom written software. All data are reported as mean ± standard error. Sample sizes were not predetermined using statistical methods, however all sample sizes were similar to previously reported behavioral, molecular, and *in vivo* electrophysiological studies^37, 71^. To measure statistical significance for differences in behavior between control and defeated mice, two-way or repeated measures ANOVAs followed by Fisher’s PLSD post-hoc testing when appropriate were performed.

Two-tailed t-tests planned *a priori* were used to assess the effects of mPFC-dPAG inhibition separately in control and defeated mice. fEPSP data was analysed using a repeated two-way ANOVA. For analysis of local field potential data we used non-parametric Mann-Whitney *U*-tests as previously described to compare theta, beta, and low gamma coherence between control and defeated mice^37^.

## Author contributions

T.B.F. designed, performed and analyzed all experiments, except the retrograde tracer experiments that were designed, performed, and analyzed by L.M., the *in vitro* electrophysiology experiments that were designed, performed, and analyzed by Z.P. and T.B., the monosynaptic rabies experiment that was designed, performed, and analyzed by B.A.S., the evoked field potential experiments that were designed, performed, and analyzed by M.E.M., the Granger causality and power analyses that were carried out by Y.Z., and for some behavioral experiments and imaging that were performed and analyzed by A.K., V.V., L.G., A.H. and S.P. The AAV-*Syn*::Venus-2A-HAhM4D virus was packaged and tested by V.G and A.I. The wireless recording device was built by A.L.V. The project was conceived and the manuscript written by T.B.F and C.T.G with critical input from T.B.

## Acknowledgements

We thank F. Zonfrillo for animal husbandry, and P. Heppenstall (Mouse Biology Unit, EMBL) for providing the *Vglut2*::Cre mouse line. Funding was provided by EMBL (C.T.G., T.B.F.), the ERC Advanced Grant “Corefear” (C.T.G.), the Swiss National Science Foundation Advanced Fellows Program (T.B.F.), the Wellcome Trust/Royal Society Henry Dale Fellowship (098400/Z/12/Z) and Medical Research Council (MRC) grant MC-UP-1201/1 (T.B.), a Marie Sklodowska-Curie grant no. 659842 (Z.P.), and One hundred Talents Program of CAS and funding from Shenzhen city government (JCYJ20140901003938992, KQCX2015033117354153) (Y.Z.).

---

**Supplementary Figure 1.**
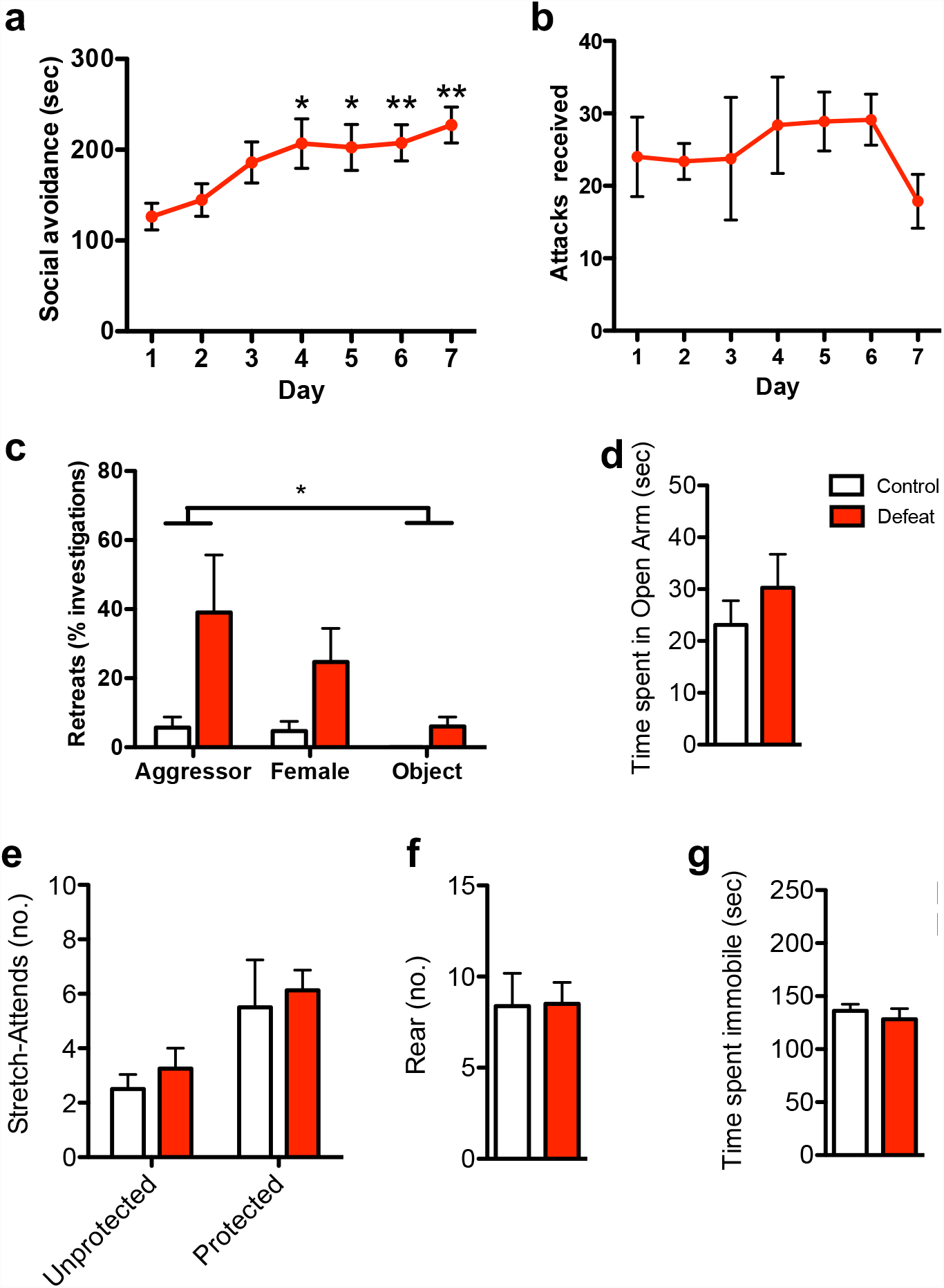
(**a**) Increased social avoidance across social defeat sessions during the anticipatory period in which the intruder was prevented from attacking the resident by a wire mesh enclosure (day: F_[6,8]_ = 5.81, p = 0.0001). (**b**) Number of attacks by the aggressor was not significantly changed across social defeat sessions. (**c**) Defeated mice made more retreats when compared to control mice, an effect observed in response to aggressors, still present in response to females, but absent in response to a novel object (defeat: F_[1,12]_ = 6.5, p = 0.026, stimulus: F_[2,12]_=3.48, p=0.047). (**d-f**) Defeated mice exhibited normal anxiety-like and risk assessment behavior in the elevated plus maze. They (**d**) spent similar time in the open arm (n=17-18), and performed comparable levels of (**e**) unprotected and protected stretch attends (n=8), and (**f**) rearing as control mice (n=8). (**g**) Defeated mice exhibited normal immobility in the tail suspension test (n=9).

**Supplementary Figure 2.**
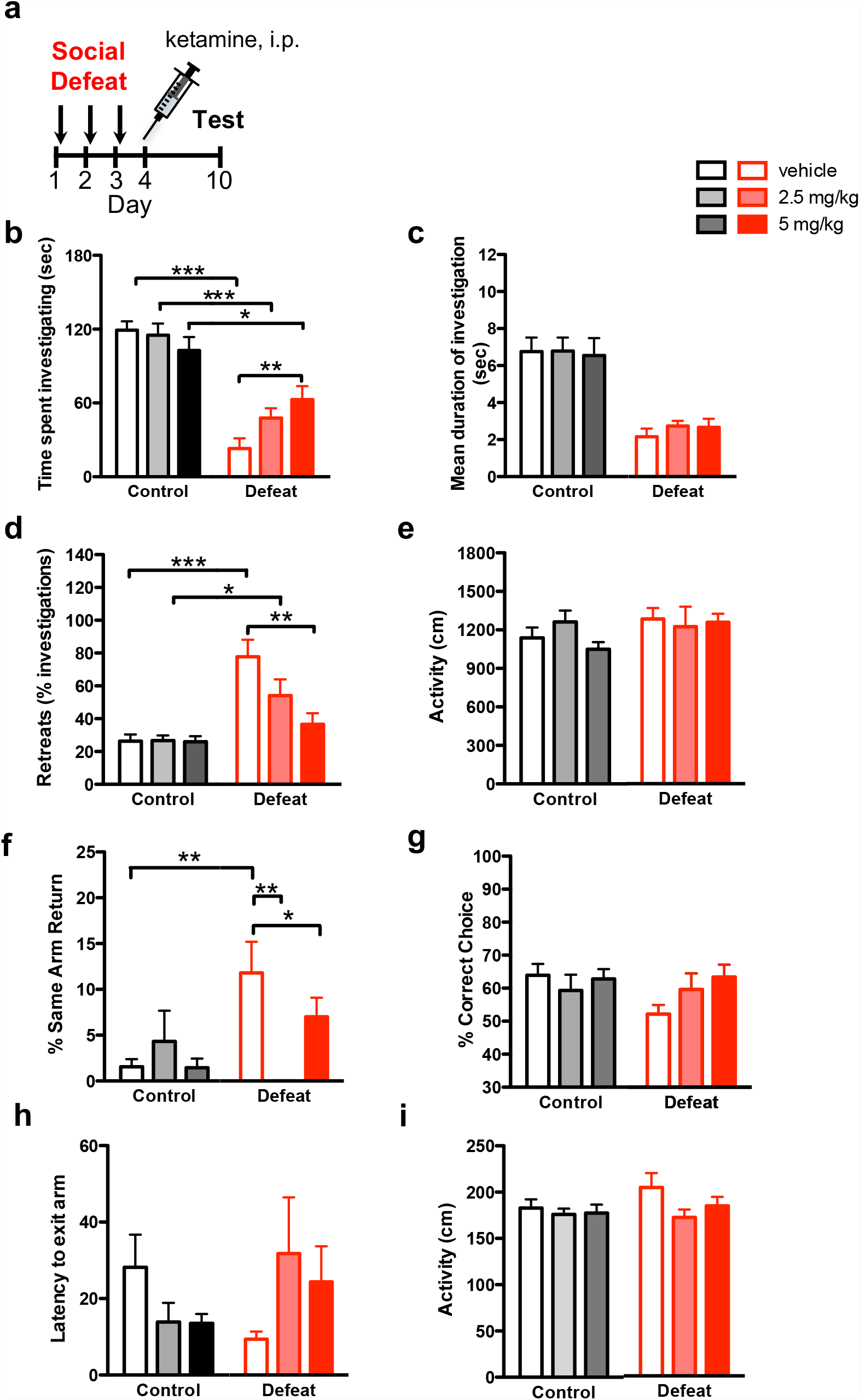
(**a**) A single dose of ketamine (2.5 and 5 mg/kg, i.p.) administered one day after the final social defeat elicited a reversal of social avoidance behavior. Socially defeated, but not control mice treated with ketamine showed a dose-dependent increase in (**b**) time investigating (defeat: F_[1, 57]_ = 65.8, p < 0.0001; defeat x ketamine: F_[2,57]_ = 4.3, p = 0.018), no change in (**c**) duration of time spent investigating, and decrease in (**d**) retreats (defeat: F_[1,56]_ = 31.9, p < 0.0001; defeat x treatment: F_[2, 56]_= 5.9, P=0.0048; ^+^p < 0.1, *p < 0.05; **p < 0.01; ***p < 0.001) (e) No effect of ketamine on baseline home cage locomotor behavior was observed. (**f, g**) Defeat-induced deficits in working memory were reversed by ketamine treatment (SARs, defeat x drug: F_[2, 50]_ = 6.3, P=0.0037; *P<0.05, **P<0.01, ***P<0.001). (**h**) Latency to exit the start arm and (**i**) overall distance travelled in the Y-maze were not altered by defeat or ketamine treatment. (^+^0.05<P<0.1, *P<0.05, **P<0.01). n=9-14/treatment.

**Supplementary Figure 3.**
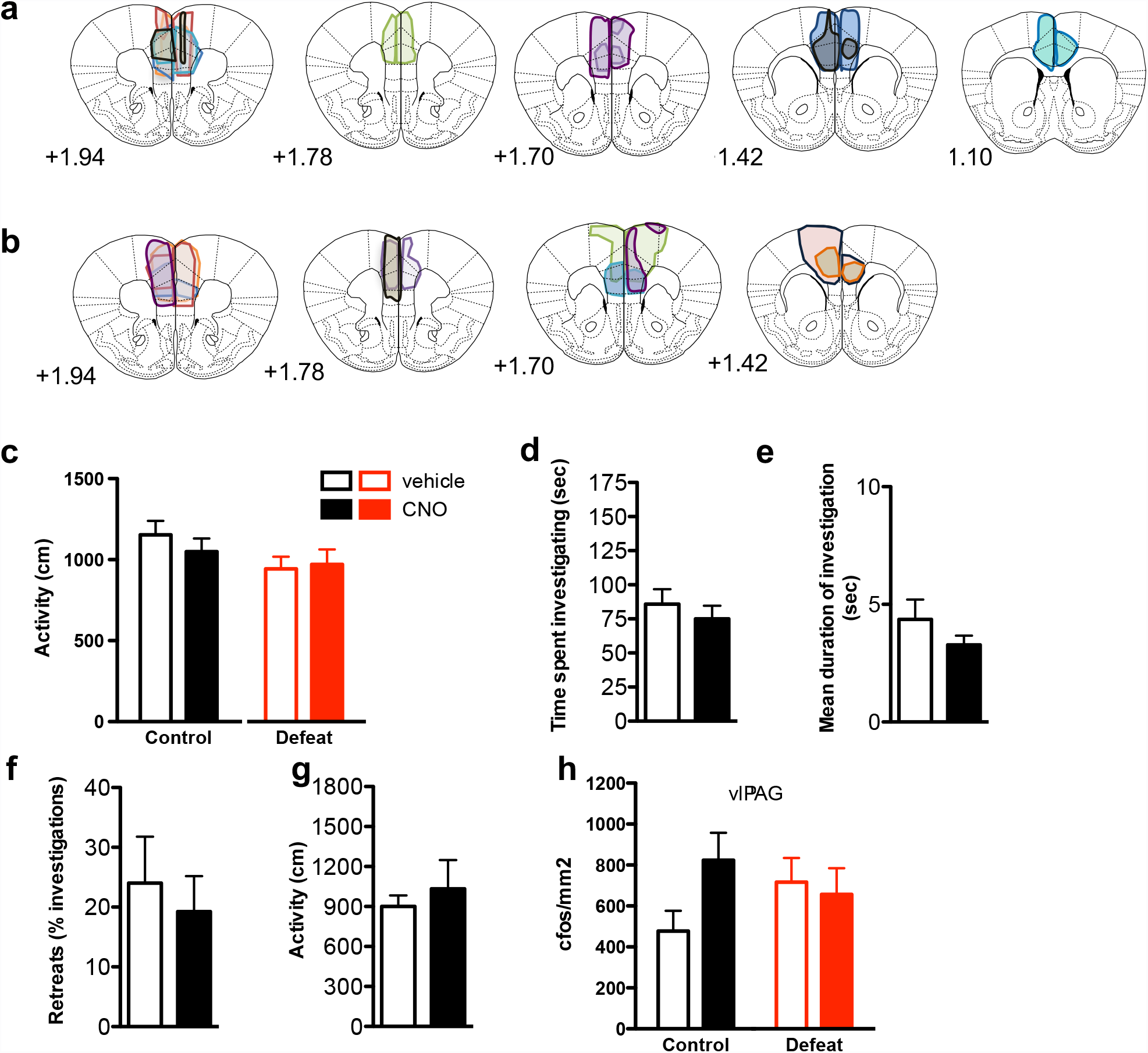
Area of viral infection (AAV-*Syn*::Venus-2A-HAhM4D-WPRE) visualized by endogenous Venus in mPFC of (**a**) control and (**b**) defeated mice administered CNO. (**c**) Distance travelled in the home cage in (left) control and (right) defeated mice after intra-PAG administration of CNO or vehicle. (**d-g**) Mice were infected bilaterally in mPFC with AAV expressing Venus fluorescent protein and HA-tagged hM4D (AAV-*Syn*::Venus-2A-HA-hM4D), implanted with a guide cannula over SuColl, subjected to control conditions, and infused locally in SuColl with CNO or vehicle before testing for social interaction. Behavior of CNO-treated control animals was indistinguishable from vehicle-treated control mice. Control mice administered CNO prior to testing displayed normal (**d**) time investigating the aggressor, (**e**) investigation bouts, (**f**) retreats and (**g**) overall activity. (**h**) Quantification of cFos immunopositive cells in ventrolateral (vl) PAG.n=6-8.

**Supplementary Figure 4.**
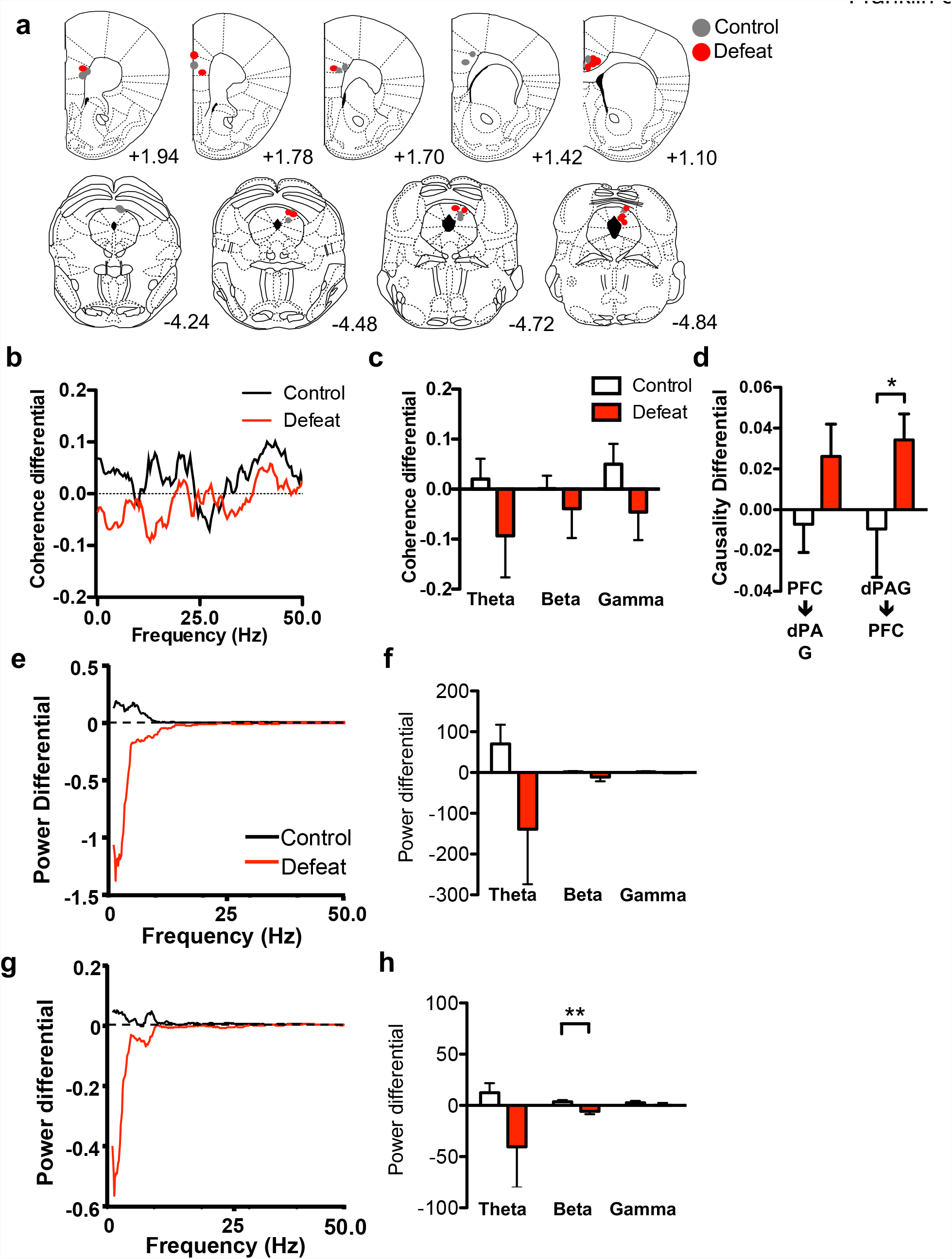
(**a**) Electrode placements for LFPs recorded in mPFC and dPAG of control (grey) and defeated (red) mice. (**b**, **c**) Relative coherence (coherence differential) was not significantly different in defeated mice compared to control animals when distal to the aggressor. (**d**) Relative causality (causality differential) was significantly higher in the dPAG->mPFC direction in defeated mice compared to control animals when mice were distal to the aggressor (U=8, p=0.038). Power spectra differential between day 1 and day 3 in (**e**, **f**) mPFC and (**g**, **h**) AG when control and defeated mice were distal to the aggressor. Defeated mice had lower power in the beta band in the PAG compared to control mice (U=6, p=0.0047). Power spectra were averaged across mice. Power in each frequency band was calculated as the sum of the power values. n=7-8, *P<0.05.

**Supplementary Figure 5.**
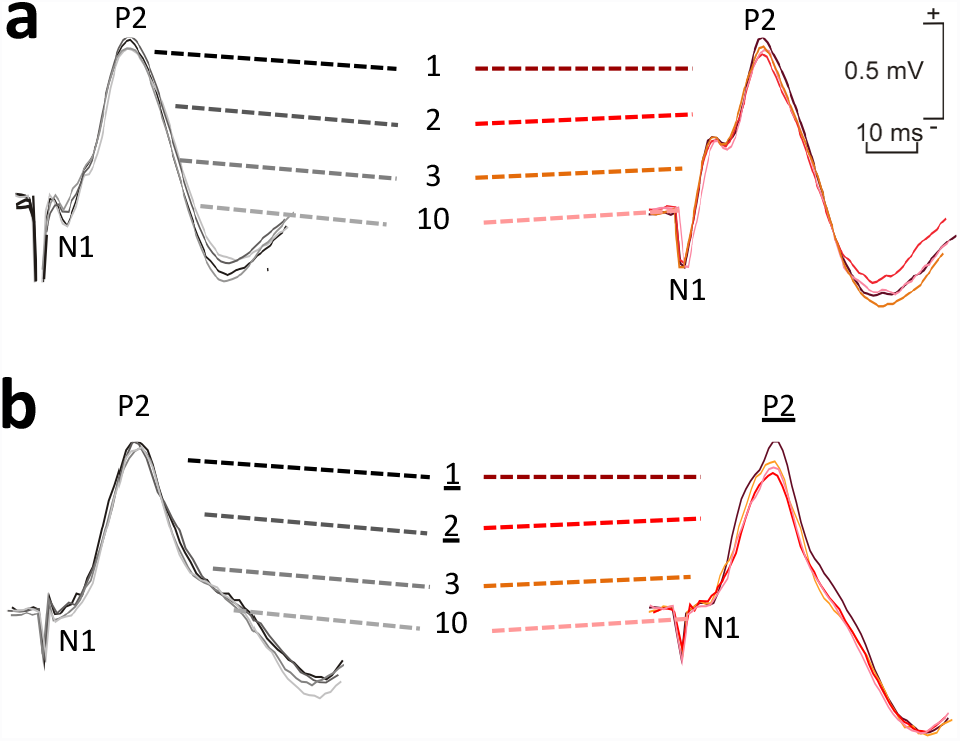
Superimposed recordings illustrating extracellular synaptic **field** potentials recorded at (a) dPAG following electrical stimulation of mPFC and at (b) mPFC following electrical stimulation of MDT along the different sessions. (gray scale: sensory group; red scale: defeated group).

**Supplementary Figure 6.**
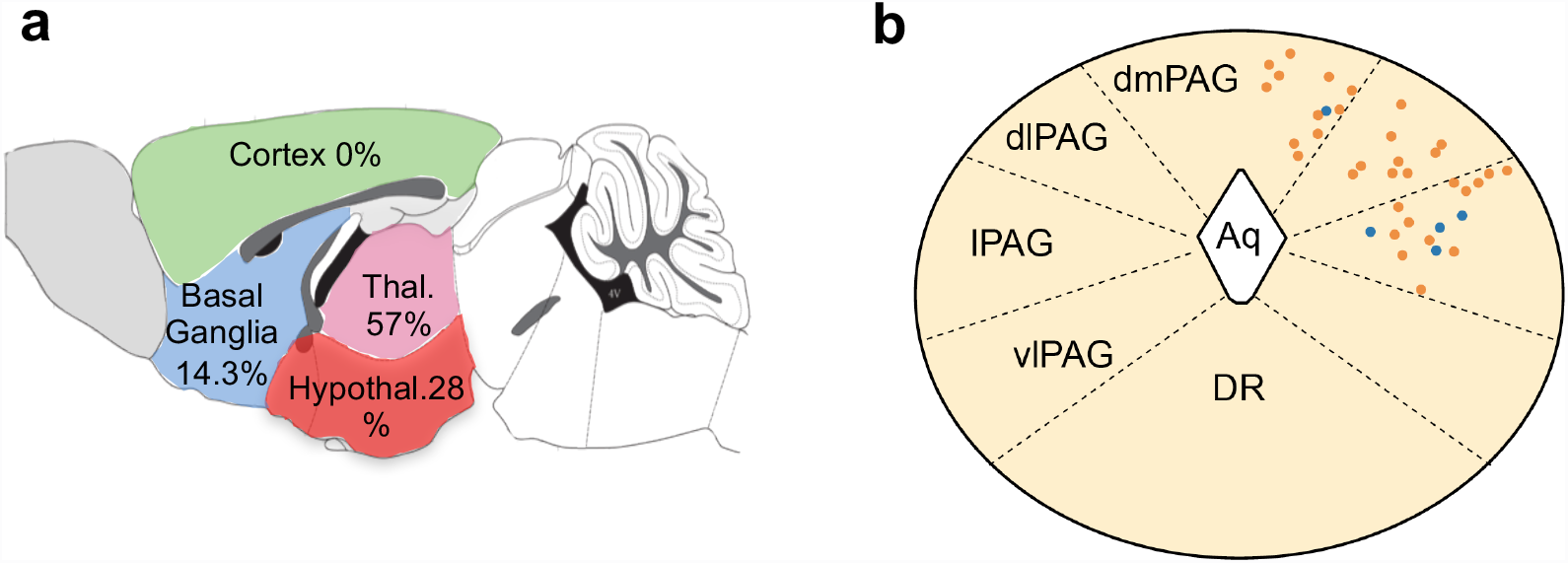
(**a**) Schematic representation of rabies-infected neurons (GFP+. mCherry-) in the telencephalon and diencephalon of *Gad2*::Cre animals infected in dPAG. No neurons were found in the olfactory bulb, hippocampus, or cortex (grey). Areas not counted (midbrain and hindbrain) are indicated in white. (**b**) Schematic showing the location of all Vglut2+ neurons from which whole-cell recordings were made. Blue circles represent neurons with monosynaptic inputs from the PFC, and orange circles are neurons without PFC input. Aq – cerebral aqueduct; dmPAG – dorsomedial PAG; dlPAG – dorsolateral PAG; lPAG – lateral PAG; vlPAG – ventrolateral PAG; DR – dorsal raphe.

**Supplementary Figure 7.**
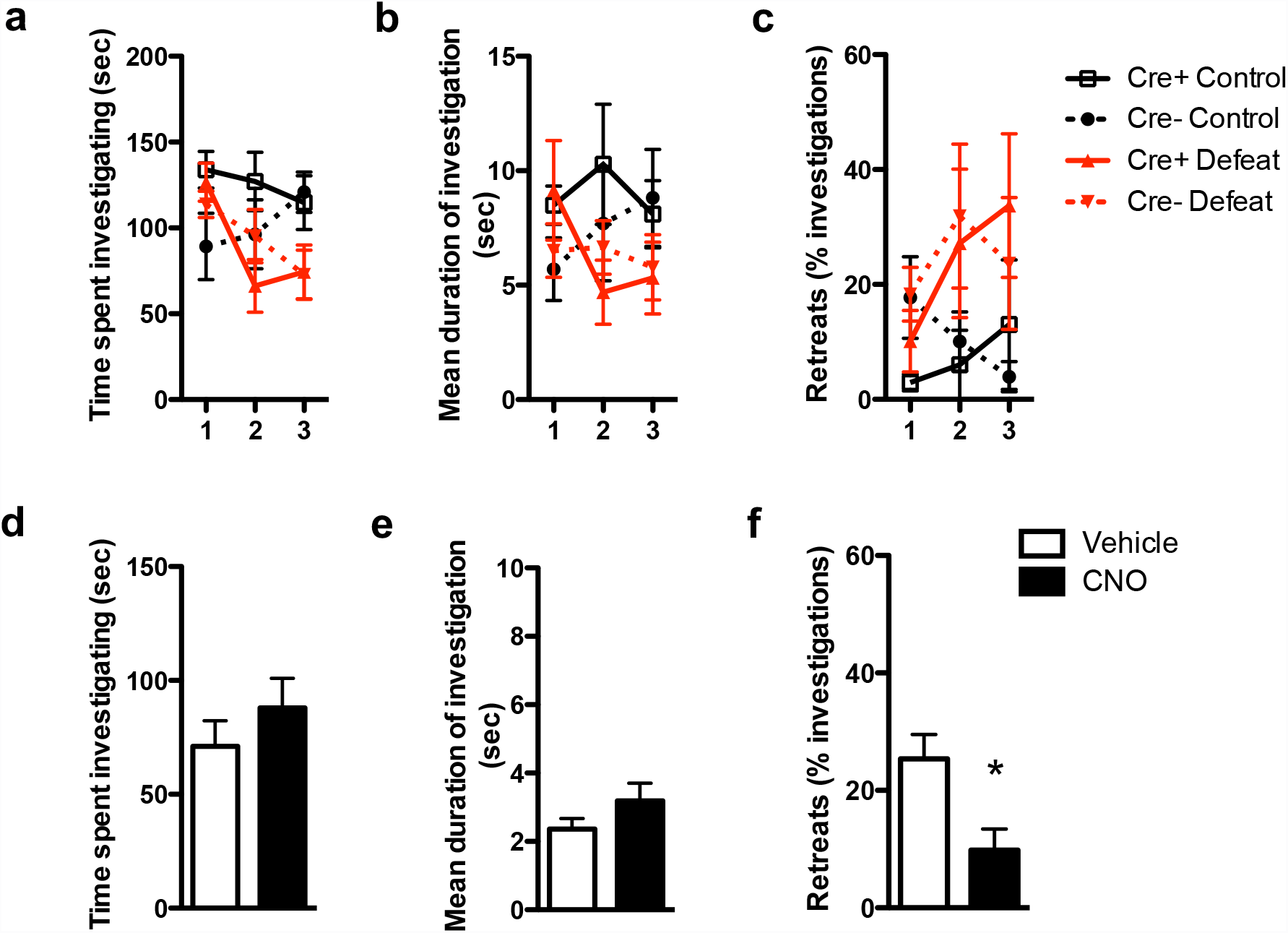
**(a-c)** Cre+ and Cre-mice behaved similarly during the three days of acquisition under both control and defeat conditions. Selective hM4D-mediated inhibition of GAD2+ neurons in dPAG. (**d-e**) *GAD2*::Cre mice were infected with AAV-*Syn*::DIO-hM4D-mCherry in the dPAG, subjected to control conditions, and treated with vehicle or CNO prior to social interaction testing. Systemic administration of CNO had (**d**) no effect on time spent investigating or (**e**) mean duration of investigation, but (**f**) reduced the percentage of retreats compared to vehicle-treated animals (t_(11)_=2.2, p=0.016). n=6-7.

**Supplementary Table 1.**

|  | CTB647-labeled neurons | GABAergic neurons | GABAergic CTB647-labeled neurons |
| --- | --- | --- | --- |
| Anterior cingulate | 130 | 1442 | 0 |
| Prelimbic | 259 | 1108 | 0 |
| Infralimbic | 194 | 976 | 0 |
| Total | 583 | 3526 | 0 |

**Supplementary Table 2.**

|  | Mouse |  |  |  |  |  |  |  | Average weighted<br>to no. of starter cells | Total no.<br>of cells |
| --- | --- | --- | --- | --- | --- | --- | --- | --- | --- | --- |
|  | #1 | #2 | #3 | #4 | #5 | #6 | #7 | #8 |  |  |
| **Cortex** | **2** | **0** | **7** | **0** | **76** | **27** | **7** | **63** | **28.0** | **182** |
| FrA |  |  |  |  |  |  |  | 1 | 0.2 | 1 |
| M2 |  |  | 1 |  |  |  |  | 1 | 0.3 | 2 |
| S1 |  |  | 1 |  | 20 | 10 |  | 4 | 4.2 | 35 |
| S2 |  |  |  |  | 3 |  | 1 | 1 | 0.8 | 5 |
| Au |  |  |  |  | 6 | 1 |  | 9 | 3.0 | 16 |
| V1 |  |  |  |  | 1 |  |  | 10 | 2.5 | 11 |
| V2 |  |  |  |  | 17 | 6 |  | 9 | 4.7 | 32 |
| mPFC | 2 |  |  |  | 8 | 5 | 3 | 2 | 2.4 | 20 |
| VO |  |  |  |  |  |  | 2 | 1 | 0.6 | 3 |
| LO |  |  |  |  | 2 |  |  | 2 | 0.7 | 4 |
| Ai |  |  | 1 |  | 2 |  |  |  | 0.3 | 3 |
| Ect |  |  |  |  |  |  |  | 1 | 0.2 | 1 |
| RSA |  |  |  |  | 7 | 2 | 1 | 19 | 5.7 | 29 |
| RSG |  |  |  |  | 6 |  |  | 3 | 1.5 | 9 |
| LPtA |  |  | 4 |  | 4 | 3 |  |  | 1.0 | 11 |
| **Striatal-like Structures** | **1** | **0** | **0** | **0** | **9** | **40** | **1** | **5** | **5.3** | **56** |
| LS | 1 |  |  |  | 4 | 2 | 1 |  | 0.8 | 8 |
| MeA |  |  |  |  | 1 |  |  | 1 | 0.4 | 2 |
| BMA |  |  |  |  | 1 |  |  |  | 0.1 | 1 |
| CeA |  |  |  |  | 3 | 38 |  | 4 | 4.0 | 45 |
| **Pallidum-like Structures** | **0** | **0** | **0** | **0** | **1** | **39** | **0** | **6** | **4.2** | **46** |
| BSTS |  |  |  |  | 1 | 36 |  | 2 | 3.1 | 39 |
| VP |  |  |  |  |  | 1 |  |  | 0.1 | 1 |
| SI |  |  |  |  |  | 2 |  | 4 | 1.1 | 6 |
| **Thalamus** | **13** | **3** | **17** | **14** | **749** | **186** | **0** | **446** | **218.8** | **1428** |
| LGN | 13 | 2 | 10 | 11 | 494 | 55 |  | 332 | 149.0 | 917 |
| AAD |  |  |  |  | 1 |  |  |  | 0.1 | 1 |
| Re |  |  |  | 1 | 1 | 3 |  | 14 | 3.9 | 19 |
| LDVL |  |  |  |  | 1 |  |  |  | 0.1 | 1 |
| RT |  |  |  |  | 5 | 4 |  | 9 | 3.0 | 18 |
| LPRM |  |  |  |  | 2 |  |  |  | 0.3 | 2 |
| SPF |  |  |  |  | 83 |  |  | 33 | 18.3 | 116 |
| APTD |  |  |  |  | 33 | 6 |  |  | 4.6 | 39 |
| PSTh |  |  |  |  | 7 | 59 |  | 11 | 7.6 | 77 |
| PoT |  |  |  |  | 68 | 31 |  |  | 10.8 | 99 |
| LPL |  |  |  |  |  |  |  | 39 | 9.2 | 39 |
| Undefined thalamus |  | 1 | 7 | 2 | 54 | 28 |  | 8 | 11.8 | 100 |
| **Hypothalamus** | **7** | **3** | **34** | **19** | **500** | **426** | **8** | **522** | **225.4** | **1519** |
| Pe |  |  |  |  |  | 1 |  |  | 0.1 | 1 |
| LPO |  |  | 1 |  |  | 2 |  |  | 0.2 | 3 |
| DM |  |  | 2 |  | 11 | 12 | 1 | 27 | 8.9 | 53 |
| AH |  |  |  |  | 37 | 15 |  | 37 | 14.5 | 89 |
| MPA |  |  | 4 | 1 | 7 | 28 |  | 3 | 4.1 | 43 |
| MPO |  |  | 2 |  | 5 | 25 | 2 | 18 | 7.1 | 52 |
| PMD | 1 |  |  | 1 | 36 | 11 |  |  | 5.7 | 49 |
| PaP |  |  |  |  |  | 1 |  | 8 | 2.0 | 9 |
| PVN |  |  | 1 |  | 4 |  |  |  | 0.6 | 5 |
| VMHa |  |  |  |  | 14 | 7 | 2 | 9 | 4.7 | 32 |
| VMHdm | 1 | 1 | 9 | 2 | 42 | 19 | 2 | 52 | 20.5 | 128 |
| VMHvl |  | 1 |  |  | 5 | 15 |  | 12 | 4.5 | 33 |
| PH |  |  |  |  | 24 | 34 |  |  | 5.4 | 58 |
| PeF |  |  |  |  | 4 | 31 |  |  | 2.7 | 35 |
| LH | 1 |  |  | 6 | 36 | 98 |  | 86 | 33.4 | 227 |
| LPO |  |  | 1 |  |  | 2 |  |  | 0.2 | 3 |
| Tu |  |  |  |  | 2 |  |  |  | 0.3 | 2 |
| TC |  |  |  |  |  | 11 |  |  | 0.8 | 11 |
| ZI | 4 | 1 | 14 | 9 | 273 | 114 | 1 | 270 | 110.0 | 686 |
| **density of starter cells in**<br>**dPAG (n/mm<sup>2</sup>)** | **21.6** | **38.6** | **106.4** | **468.8** | **203.6** | **110.3** | **276.8** | **375.5** |  |  |

**Supplementary Table 3.**

|  | Mouse |  |  |  |  |  |  | Average weighted to no.<br>of starter cells | Total no.<br>of cells |
| --- | --- | --- | --- | --- | --- | --- | --- | --- | --- |
|  | #1 | #2 | #3 | #4 | #5 | #6 | #7 |  |  |
| **Cortex** | 0 | 0 | 0 | 0 | 0 | 0 | 0 | 0.0 | 0 |
| **Striatal-like Structures** | 0 | 0 | 0 | 0 | 0 | 1 | 1 | 0.5 | 2 |
| LS |  |  |  |  |  | 1 |  | 0.3 | 1 |
| CeA |  |  |  |  |  |  | 1 | 0.2 | 1 |
| **Pallidum-like Structures** | 0 | 0 | 0 | 0 | 0 | 0 | 0 | 0.0 | 0 |
| **Thalamus** | 3 | 1 | 0 | 2 | 2 | 0 | 0 | 1.0 | 8 |
| LGN | 3 | 1 |  | 1 | 2 |  |  | 0.9 | 7 |
| Undefined thalamus |  |  |  | 1 |  |  |  | 0.1 | 1 |
| **Hypothalamus** | 0 | 0 | 0 | 2 | 2 | 0 | 0 | 0.6 | 4 |
| MPA |  |  |  |  | 1 |  |  | 0.2 | 1 |
| PH |  |  |  | 1 |  |  |  | 0.1 | 1 |
| ZI |  |  |  | 1 | 1 |  |  | 0.3 | 2 |
| **density of starter cells in**<br>**dPAG (n/mm<sup>2</sup>)** | 70.2 | 59.8 | 21.5 | 82.1 | 94.9 | 172.0 | 103.7 |  |  |

